# Counting-based inference of mutant growth rates from pooled sequencing across growth regimes

**DOI:** 10.1101/2025.10.10.681719

**Authors:** Deniz Sezer, Erdal Toprak

## Abstract

Time-resolved sequencing of pooled mutants is widely used to track their frequencies under selection pressure, thereby revealing variants that are enriched or depleted. Here, we address how to quantify variant growth rates by analyzing the temporal dimension of the counts data through a model of growth. For exponential growth, we first study weighted least-squares fitting and show that non-linear fitting based on the *softmax* transformation exhibits more favorable properties than the currently employed linear regression. We then argue that direct maximization of the likelihood of the noise model should be preferred over least-squares fitting. For a multinomial model of counting noise, we adopt variational Bayesian inference to additionally quantify uncertainties in the estimated growth rates. We provide closed-form expressions for the experimentally practical case of sequencing only at the beginning and at the end of the experiment. Finally, we extend maximum-likelihood estimation and variational Bayesian inference to logistic and Gompertz growth, which serve as illustrative examples of general, non-exponential growth models formulated in terms of a small number of parameters per variant. The ability to incorporate arbitrary growth models within the developed inference framework opens new opportunities for high-throughput estimation of diverse biochemical parameters that influence growth.

**Author summary:** Simultaneously tracking the relative abundances of thousands of genetically distinct cellular variants over time is now possible through deep sequencing and other counting-based methods. This capability provides a quantitative window into increasingly broad regions of the combinatorial fitness landscape—well beyond those explored by natural evolution. While identifying a few engineered variants that survive under extreme selection is valuable, quantitative mapping of the fitness landscape requires accurate inference of growth rates for the entire variant population. Here, we revisit three approaches for estimating growth rates from sequencing count data: least-squares fitting, maximum likelihood estimation, and variational Bayesian inference. By clearly delineating the probabilistic model of counting noise and the deterministic model of variant growth—both jointly required for inferring growth rates from time-resolved sequencing counts—we show how exponential growth can be replaced by any alternative growth model. Applying the developed analysis framework to models in which growth rates are expressed in terms of microscopic biochemical parameters will enable the high-throughput inference of fundamental kinetic and biophysical constants from sequencing data.

## 1 Introduction

Recent advances in deep sequencing and gene synthesis technologies have enabled the simultaneous interrogation of thousands of genetically distinct pooled variants growing under nearly identical conditions within the same culture [1]. Depending on the experimental design, these variants may represent distinct gene knockouts [2] or knockdowns [3] of a parental genome, or they may differ in the sequence of one [4–6], two [7], or multiple genes whose products influence growth under the studied conditions.

On a qualitative level, screening thousands of variants in parallel increases the chance of identifying a few exceptionally fit mutants among a combinatorially vast space of possible genotypes [6]. Quantitatively, estimating the fitness of each variant from changes in its relative abundance over time—as measured through sequencing-based counting—provides a systematic framework for mapping epistatic interactions between mutations. Elucidating the mechanistic basis of these interactions, in turn, offers deeper insights into the underlying biological processes.

This work focuses on the quantitative estimation of variant growth rates from sequencing-based counts. Specifically, we consider growth *competition* experiments in which a diverse population of pre-existing variants is propagated in parallel for several generations—a time scale short enough that the spontaneous occurrence of new mutations can be neglected. This setup contrasts with *evolution* experiments, which typically span hundreds of generations and are designed to study the emergence and fixation of novel genetic changes [8].

Estimating growth rates requires sequencing at least two time points along the competition trajectory, and many studies sequence only at the beginning and end of the experiment [5]. With two points only, assuming exponential growth between them, the growth rate can be inferred by fitting a line through two log-transformed abundance values. When sequencing data are collected at more than two time points, the question arises of how best to integrate the additional temporal information. One intuitive approach, implemented for example in the Enrich2 package for analyzing deep mutational scanning data [9], is to fit a weighted least-squares line through the time points. In contrast, likelihood-based approaches model sequencing counts at each time point through either a Poisson [8], negative-binomial [10], or multinomial [11] distribution, and use exponential growth to link the parameters of the distribution across time. More recently, this latter approach has been extended to Bayesian inference of growth rates, where uncertainties of the estimates are an integral part of the statistical model [11].

Because all these analytical approaches assume sustained exponential growth, maintaining such conditions experimentally is essential. This can be achieved using a turbidostat [3, 7], which continuously dilutes the culture to maintain constant cell density, or through frequent manual dilutions [12]. In fixed-duration batch protocols, however, nutrient depletion and waste accumulation progressively limit growth, making the assumption of uninterrupted exponential behavior—including between dilutions—increasingly unrealistic.

More generally, a central assumption in estimating growth rates from pooled competition assays is that variants do not interact with one another, despite sharing the same physical environment. This stands in contrasts to selection experiments on microbial communities, where interspecies interactions that modulate individual growth rates are essential for determining both the resilience and long-term composition of the community [13–15]. Similar inter-variant interactions may also occur in pooled competition assays. A well-studied example arises in the context of beta-lactam antibiotics: resistant bacteria expressing beta-lactamase enzymes hydrolyze the antibiotic, lowering its concentration in the medium and thereby reducing selection pressure for all members of the mixed population, including the susceptible ones [16, 17]. Because of this complication, a recent study from our group on the beta-lactamase TEM-1 used the area under the variant growth curve as a proxy for fitness, rather than calculating explicit growth rates [6].

Here, as a first step toward accommodating potential inter-variant interactions in pooled competition assays, we extend the sequencing-based analysis of growth rates to growth models with saturation, thereby moving beyond the assumption of variant independence and exponential growth. Unlike the specific interactions central to studies of microbial communities, the reduction of growth near saturation represents a non-specific interaction affecting all variants. To distinguish the initial, near-exponential phase from the subsequent saturation regime, sequencing at more than two time points is required. We therefore revisit existing strategies—least-squares fitting and time-parametrized likelihood methods—for analyzing time-resolved sequencing data.

When modeling the noise in the experimental counts, we focus on counting noise arising from the partitioning of a finite number of sequencing reads among the variants. Other sources of variability—such as noise from the PCR amplification or physiological heterogeneity among isogenic variants—are inevitable [8], but are not considered in our analysis. The multinomial model of counting noise that we adopt, respects the compositional constraint that variant fractions sum to one [18]. Its characteristic negative covariances capture the fact that an increase in the fraction of one variant necessarily entails a decrease in the fractions of the others. This interdependence is lost when the counting noise of individual variants is modeled independently with Poisson [8] or negative-binomial [10] distributions.

In contrast to previous studies, we express the variant fractions as a *softmax* transformation of the logarithms of the variant abundances [Eq (13)]. Under exponential growth, this parametrization yields an exact, closed-form expression for the time-dependence of the variant fractions [Eq (38)]. Without this transformation, the dynamics of the fractions cannot be integrated directly—even in exponential growth—because they depend explicitly on the mean population growth rate [8, 10–12]. As a result, likelihood-based methods [8, 10, 12], including the recent Bayesian formulation [11], face the problem of estimating the mean growth rate as a prerequisite to estimating the variant-specific growth rates [8, 10–12]. Our parametrization removes explicit reference to the mean population growth rate, thereby eliminating the need to estimate it as an intermediate step.

We model variant growth deterministically. Because the cell is the minimal replicating unit, we assume that each cell can be assigned to a single genetic variant, with cells sharing identical genotypes belonging to the same variant. Situations that complicate this simple correspondence—such as ongoing genome replication without cell division, or plasmid-borne variants with variable copy numbers—should be analyzed on a case-by-case basis.

When modeling saturation, we integrate the growth dynamics numerically for given parameter values of the growth model, and use automatic differentiation [19] to compute gradients of the relevant loss function—likelihood or evidence lower bound—with respect to these parameters. Although only the sequencing time points ultimately contribute to the loss function and its gradients, the numerical integration step size is independent of the temporal resolution of the sequencing data and can be chosen to be much smaller. This finer integration effectively interpolates the growth trajectories between sequencing time points, enabling potential comparisons with complementary high-resolution measurements such as optical density or metabolomics data. If available, such data could serve as additional constraints for parameter estimation in the growth model. (This possibility is not explored in the present paper.)

Given the current costs of deep sequencing, the sampling frequency along the time axis is likely to remain limited for the near future. While a relatively small number of sequencing time points is sufficient for estimating growth rates in the absence of direct inter-variant interactions, it does not permit the simultaneous estimation of interaction parameters, whose number can be very large. In the case of pairwise interactions, for example, the number of parameters scales quadratically with the number of variants, becoming prohibitively large even for tens of variants. For this reason, the estimation of such specific pairwise interactions is not attempted in the present work.

In Sec 2, we introduce the necessary prerequisites. In addition to summarizing known concepts to make the paper self-contained for a broad audience, this section contains two novel insights. First, we connect the probabilistic model of counting noise and the deterministic model of growth through the *softmax* transformation (Sec 2.3). This mathematical relationship clarifies why only relative growth rates can be inferred from sequencing data in the case of exponential growth, and how saturating growth provides access to absolute growth rates. Second, for the multinomial probabilistic model of counting noise (Sec 2.1), we compare exact Bayesian analysis with Dirichlet posterior (Sec 2.2) and variational Bayesian inference with Gaussian posterior (Sec 2.4). Because the latter posterior introduces twice as many parameters compared to the former, we study how variational inference obtains values for 2*K* parameters from only *K* observed numbers.

Our results are presented in Sec 3, where we address the question of how to integrate the temporal dimension of the sequencing data through the model of growth. Specifically, in Sec 3.1 we examine the weighted least-squares procedure implemented in the analysis tool Enrich2 [9], and propose an alternative mean-squared error with more favorable fitting properties. Then, in Sec 3.2 we reformulate the problem as maximum-likelihood estimation, eliminating the need for least-squares fitting. We also revisit the estimation of growth rates in the special case of sequencing at only the beginning and end of the experiment. Subsequently, in Sec 3.3 we adopt a variational-inference framework [11] to assign uncertainties to the estimated growth rates, and derive new Bayesian estimators for sequencing at only two time points. While these first section focus on exponential growth, the developed maximum-likelihood and variational-inference approaches are applied in Sec 3.4 to two models with saturation—logistic and Gompertz growth [20].

Our conclusions are summarized in Sec. 4, and additional details are provided in the Supporting Information (hereafter referred to as SI).

## 2 Methods

### 2.1 Multinomial model of counting noise

We denote the number of sequencing reads assigned to variant *k* at time point *t*_*τ*_ by *n*_*kτ*_ . The total number of reads across all *K* variants at *t*_*τ*_ is then

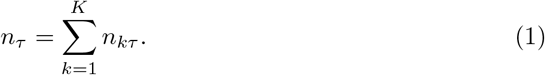

There are *T* sequencing time points in total, and we place the origin of the time axis at *t*_0_ = 0, thus *τ* = 0, 1, …, *T* ∗ 1. We use *K*-dimensional vectors, indicated by a lowercase bold symbol, to collectively refer to properties of all variants at a single time point, e.g., **n**_*τ*_ = (*n*_1*τ*_, …, *n*_*Kτ*_). The complete experimental dataset of *K × T* counts can be viewed as the matrix **N** = [*n*_*kτ*_], whose *τ* th column is **n**_*τ*_ .

We model the experimental read counts at a given sequencing time point as random draws from a multinomial distribution parametrized by the true variant fractions in the culture. With *N*_*k*_(*t*) denoting the number of cells that correspond to variant *k* at time *t*, the fraction of this variant is

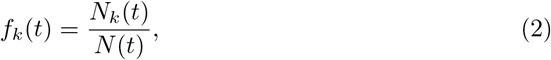

where the normalization is by the total number of cells

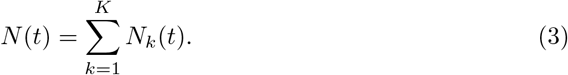

The multinomial probability of observing **n**_*τ*_ sequencing counts, given the true fractions **f**_*τ*_ = **f** (*t*_*τ*_), is

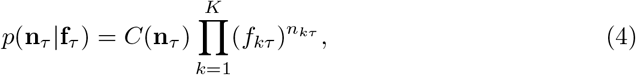

where the normalization factor *C*(·) is not of direct relevance here, since it does not depend on the fractions.

To estimate the parameters **f**_*τ*_ from a given realization of the counts **n**_*τ*_, we maximize the probability (4), now regarded as a function of the variant fractions and referred to as *likelihood*. Retaining only terms that depend on **f**_*τ*_, the natural logarithm of the likelihood is

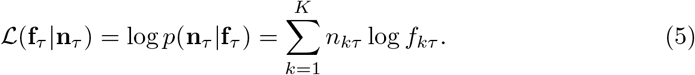

Maximizing this log likelihood, subject to the constraint that the fractions of all variants sum to one, yields the maximum-likelihood (ML) estimates (see SI, Sec S2.1.1)

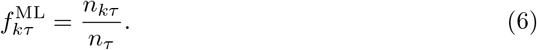

Thus, the simple ratio of the counts of variant *k* to the total variant counts, which one would compute even without formal statistical analysis, is exactly the maximum-likelihood estimate of the true fraction of that variant. This outcome is, in a sense, inevitable, as estimating *K* variant fractions from *K* observed counts leaves little room for alternative estimators. Because each observed count informs exactly one parameter, the estimates (6) do not benefit from any averaging over the stochastic fluctuations inherent in the counts.

### 2.2 Bayesian inference of the variant fractions

Maximizing the likelihood yields point estimates of the parameters of the multinomial distribution. In Bayesian inference, the parameters **f**_*τ*_ are promoted to random variables that have their own probability distribution, just like the counts **n**_*τ*_ . Unlike the counts, however, the variant fractions are not observed in the experiment and remain hidden, or *latent*. The objective is to determine *p*(**f**_*τ*_| **n**_*τ*_), the *posterior* probability distribution of these latent variables given the observed data.

The tool leveraged to this end is the Bayes theorem [21]

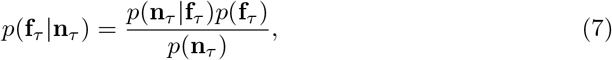

where the generative model of the data, *p*(**n**_*τ*_ |**f**_*τ*_), and the *prior* distribution of the parameters, *p*(**f**_*τ*_), are treated as known inputs, while the posterior is the desired output. (The denominator ensures that the posterior is properly normalized over the fractions.)

For the multinomial generative model (4), a Dirichlet prior yields a posterior that is also Dirichlet but with different parameters. If 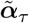 are the parameters of the prior and ***α***_*τ*_ those of the posterior, then (see SI, Sec S2.2)

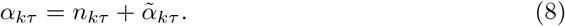

In other words, for each variant, the observed count *n*_*kτ*_ is increased by the prior parameter 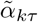 to produce the posterior parameter *α*_*kτ*_ . The prior parameters therefore act as pseudo-counts that are added to the actual counts.

Because variant counts generally differ across sequencing time points, the posterior parameters also vary with time. However, the same prior parameters can be used for all time points, i.e., 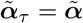. In practice, it is also common to use the same prior value for all variants, i.e., 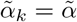. As an example, uniform Dirichlet prior corresponds to 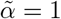, while 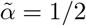, known as Jeffreys prior, is used in Enrich2 [9].

According to (8), the *K* parameters of the Dirichlet posterior are determined directly from the *K* experimental read counts at a given time point. Hence, each posterior parameter *α*_*kτ*_ is informed by a single observed count *n*_*kτ*_, just as in the maximum-likelihood estimates (6). The key difference is that maximum likelihood produced a single point estimate for the fraction of each variant, whereas the Dirichlet posterior defines a full probability distribution. This distribution can be used not only to compute the mean of each variant fraction but also to quantify the spread around the mean, for example, in the form of variance.

For the Dirichlet posterior, the means and variances of the fractions are

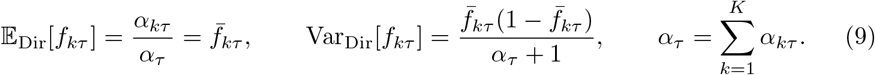

Here, 𝔼 [·] and Var[·] are operators that take the expectation and variance of their arguments, respectively, and the subscript ‘Dir’ is a reminder that these are calculated for a Dirichlet posterior.

The expected fraction of variant *k* in (9) equals the ratio between the Dirichlet parameter *α*_*kτ*_ and the sum of the Dirichlet parameters of all variants, *α*_*τ*_ . For 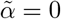 (i.e., zero pseudo-counts) this ratio becomes the maximum-likelihood estimate (6). Because of the ratio, the mean fractions remain unchanged if all posterior Dirichlet parameters are multiplied by a common constant. In contrast, the variances in (9) are sensitive to such rescaling and therefore depend on the absolute magnitude of the observed counts. Hence, *K*− 1 degrees of freedom of the Dirichlet distribution determine the mean variant fractions (which sum to one), while the remaining degree of freedom—the sum *α*_*τ*_ of all *K* Dirichlet parameters—controls the scale of the deviations from the means. Indeed, *α*_*τ*_ is often referred to as the *precision* of the Dirichlet distribution, as it reflects how tightly the distribution is concentrated around its mean [22].

For later use, we also give the means and variances

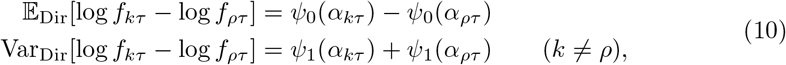

where *ψ*_0_(·) and *ψ*_1_(·) are the digamma and trigamma functions, respectively. These functions admit the asymptotic expansions [23]

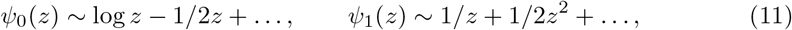

whose partial sums become increasingly more accurate as the argument *z* increases.

The transition from likelihood-based estimation of the variant fractions to Bayesian inference, outlined so far, is summarized visually on the left-hand side of Fig 1. The probabilistic model (4), where the random counts **n**_*τ*_ follow a multinomial (Mult) distribution with parameters **f**_*τ*_, is shown in the top-left panel. In this representation, random variables are depicted as circles, while the parameters of their probability distributions are indicated by points [21]. The transition to `Bayesian` modeling, in which the parameters **f**_*τ*_ are upgraded to random variables, is shown in the bottom-left panel below it. The circle of **f**_*τ*_ is left white to indicate that these random variables are latent. In contrast, shading the circle of **n**_*τ*_ in blue indicates that random realizations of these variables are available as experimental data [21]. In inference, the purpose is to estimate the values of the parameters (red points) from the data (blue-shaded circles).

**Fig 1.**
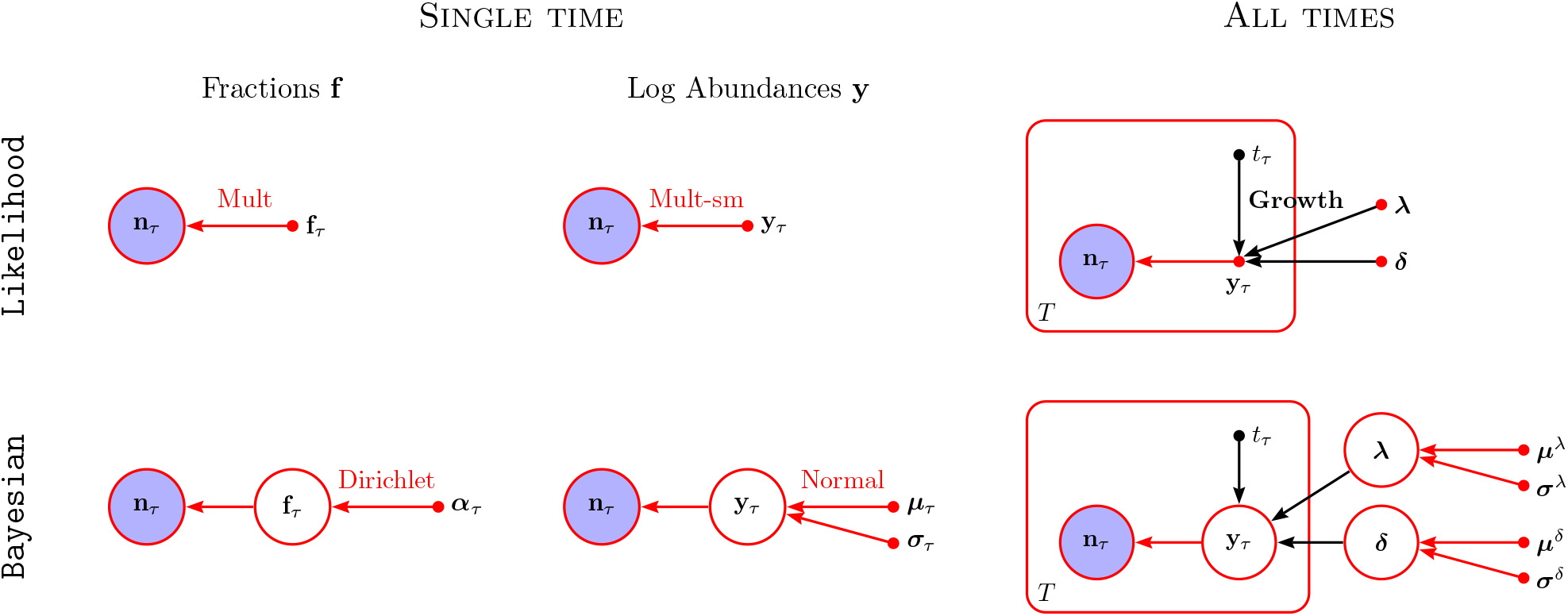
Graphical representation of the probabilistic models examined in this work. Random variables are shown with circles and parameters with points. Blue shading indicates that samples of the random variable are available as experimental data. Empty circles signify that the random variables are not observed and thus remains *hidden* or *latent*. Such latent variables arise naturally in `Bayesian` inference, where the parameters of the `likelihood` are promoted to random variables. Red arrows are used for probabilistic relationships between the variables and black arrows for deterministic functional dependence through the growth model. Variables enclosed in rectangular frame are repeated *T* times, with the index *τ* running over the repetitions. Note that only the time *t*_*τ*_ changes between the *T* repetitions, while the growth parameters ***λ*** and **δ** are shared across all time points. The task of inference consists of estimating values for the parameters represented with red points from given samples of the count variables in blue-shaded circles.

In Secs 2.3 and 2.4, we discuss the `Likelihood` and `Bayesian` models shown in the middle column of Fig 1. In the first four models, the data at time point *t*_*τ*_ is analyzed in isolation, independently from the other sequencing time points. As a result, their parameters have different values at different sequencing time points, as indicated by the subscripts *τ* . In contrast, by incorporating an explicit model of variant growth (Growth), the two rightmost models in Fig 1 parametrize the time dependence in terms of time-invariant parameters whose values can be inferred from the joint analysis of the counts across all time points. These two models are introduced in Secs 3.2 and 3.3.

### 2.3 Softmax reparametrization of the multinomial distribution

We now introduce the *log abundances*

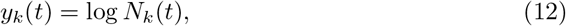

which are the natural logarithms of the variant cell numbers. From the definition of the variant fractions (2), we find

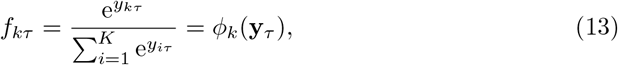

where **y**_*τ*_ = **y**(*t*_*τ*_).

The vector-valued mapping ***ϕ***(·) defined on the right-hand side of (13) is known as the *softmax* transformation [21]. (The name *logistic function* is also used in the present context [24], but it is often reserved for the special case of a single scalar variable.)

Substituting (13) into the multinomial log likelihood (5), the latter is readily expressed in terms of the log abundances of the variants as

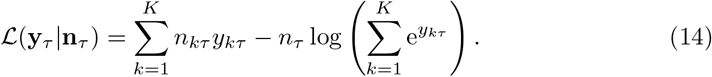

Importantly, this change of variables does not require a Jacobian factor, since we are reparametrizing the parameters of the distribution rather than the random variables themselves.

The log likelihood (14) corresponds to the middle model in the Likelihood row of Fig 1. Differentiating with respect to the log abundances *y*_*kτ*_ and zeroing the derivatives, yields the maximum-likelihood conditions (see SI, Sec 2.1.2)

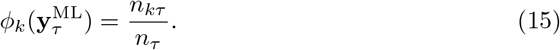

While these are equivalent to the estimates 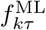 in (6), solving them for 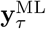 requires inverting the *softmax* function.

When working with the *softmax* transformation, it is important to recognize that it does not possess a unique inverse, since adding the same constant to the log abundances of all variants in (13) leaves the variant fractions unchanged. As a result, for experimental data reflecting only the fractions of the variants (so-called *compositional* data [18]), the log abundances can be estimated from the data only up to an arbitrary global shift. From (12), such a shift is equivalent to changing the units for *N*_*k*_(*t*), like expressing abundance in optical density (OD) instead of absolute cell counts. Nevertheless, to obtain unique numerical values of the log abundances—for instance, to perform explicit calculations or report parameter estimates—this degree of freedom must be removed by imposing an additional, yet arbitrary, constraint. Often referred to as gauge fixing [25], this step effectively amounts to selecting a particular unit for the abundances.

Given that the argument of the *softmax* function is identifiable only up to a global shift, when solving (15) for the log abundances, we can discard variant-independent additive terms. Taking the log of both sides and dropping such terms yields the maximum-likelihood estimates (see SI, Sec S2.1.2)

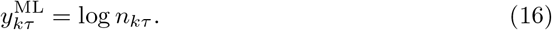

Once determined, the estimates of all variants can be shifted collectively, as desired.

In (16), the chosen gauge is left implicit. A gauge that is often selected in the literature, is to designate one variant—typically the wild type—as a reference, and then apply a global shift such that the log abundance of this reference variant is zero. In practice, this choice often arises as follows. Taking the logarithm of the *softmax* relationship in (13) yields

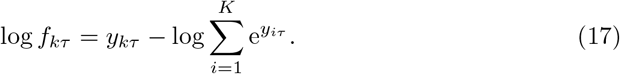

Here, the first term on the right-hand side of the equality is specific to variant *k* but the second one—due to the normalization factor in the denominator of the *softmax* transformation—is the same for all variants (although it can change at different time points *t*_*τ*_). Rather than simply dropping this term, we now cancel it by subtracting the same expression for another variant *ρ*, chosen as reference:

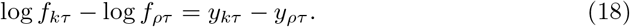

The right-hand side is now unaffected by the arbitrary global shift of the values, because it contains the difference of two log abundances.

### 2.4 Variational Bayesian inference for the log abundances

The posterior Dirichlet distribution of Sec 2.2 is for the variant fractions **f**_*τ*_ . Here, we aim to similarly obtain a posterior distribution but for the log abundances **y**_*τ*_ .

While the Dirichlet distribution is a convenient conjugate prior for the multinomial distribution (yielding a Dirichlet posterior), such conjugate prior does not exist in analytical form when the multinomial distribution is reparametrized in terms of log abundances. An approximate posterior, however, can be obtained through variational inference [11, 26]. The core idea is to restrict the posterior to a class of analytically convenient functions, and then select the parameters of the posterior so that it provides the best possible approximation to the exact but intractable posterior within that class.

The mathematical framework underlying variational inference is presented in more detail in SI, Sec S3. Here, we focus on describing the final outcome and the assumptions on which it rests. To simplify the notation, in this subsection we drop the index *τ*, with the understanding that all variables refer to the same time point.

As standard in Bayesian inference, our goal is to determine the posterior *p*(**y** | **n**). We approximate this intractable posterior within the class of distributions that factorize over the individual latent variables, a choice known as the *mean-field approximation* [21]. Further taking each factor to be Gaussian with its own mean and variance, i.e.,

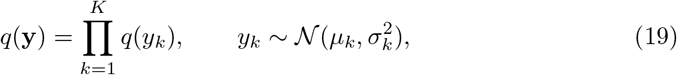

we arrive at the middle `Bayesian` model in Fig 1. Compared with the multinomial-Dirichlet model on its left, this model has twice as many free parameters—the means ***µ*** and standard deviations ***σ***—that must be inferred from the data.

For a given realization of the counts **n**, the variational parameters can be obtained by maximizing the function (see SI, Sec S3.2.3)

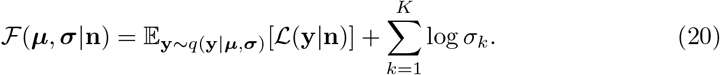

The first term on the right-hand side is the expectation of the log likelihood over the latent variables **y**, which are distributed according to the approximate posterior *q*(**y**), parametrized by ***µ*** and ***σ*** (as indicated by the subscript of the expectation operator). The second term corresponds to the cumulative entropy of the Gaussian factors forming the posterior.

In maximum-likelihood estimation, the log abundances are treated as parameters and maximizing ℒ (**y**| **n**) yields the point estimate **y**^ML^. In variational inference, by contrast, the log abundances are random variables with distribution *q*(**y**), and the parameters of this distribution are inferred by maximizing the objective (20). To see what this maximization achieves, consider what happens if the entropy term in (20) is ignored and only the expectation of the log likelihood is maximized. This can be attained by placing the means of the Gaussians at the maximum of the likelihood, i.e., ***µ*** = **y**^ML^. Since ℒ (**y**| **n**) decreases away from its maximum, the expectation is going to be maximized when the Gaussians collapse to a single point, i.e., when ***σ*** ∗ 0. Thus, the first term in (20) alone drives the approximate posterior towards the maximum-likelihood solution. The second term counteracts this collapse of *q*(**y**) by penalizing the decrease of the standard deviations. If only this second term were maximized, on the other hand, the standard deviations would grow indefinitely, pushing the approximate posterior toward a uniform distribution. Maximizing ℱ therefore strikes a balance between the uniform posterior and the maximum-likelihood solution.

For reasons explained in SI, Sec S3.1, the function ℱ is often referred to as the ELBO, short for *evidence lower bound*. It can also be viewed as a free energy obtained by adding the entropy of the posterior to the “mean energy” given by the first summand in (20) [27]. This interpretation motivates the choice of the letter ℱ.

The ELBO (20) is derived for a uniform prior, obtained in the limit of infinite standard deviations, where the terms containing the prior parameters vanish. For a non-uniform prior, the ELBO includes additional additive contributions from the prior, which ensure that maximizing everything except the expectation of the log likelihood would drive the posterior toward this non-uniform prior (see SI, Secs. S3.2.2 and S3.2.3).

When maximizing the ELBO either numerically or analytically, one must compute its gradients with respect to the variational parameters. To this end, it is convenient to express the normal random variables in (19) as

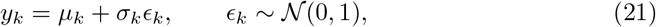

which is known as *reparametrization trick* [28, 29]. In this parametrization, the randomness of the latent variable *y*_*k*_ is carried by the *standard* normal variable *ϵ*_*k*_ with zero mean and unit variance, while *µ*_*k*_ and *σ*_*k*_ are deterministic parameters. The expectation over **y** in (20) can now be replaced by expectation over ***ϵ***, which we denote by 𝔼_*ϵ*_[·]. Note that 𝔼_*ϵ*_[*ϵ*_*k*_] = 0.

So far, we have not used the functional form of the generative model. Substituting the log likelihood (14) of the multinomial-*softmax* distribution into the ELBO (20), we obtain

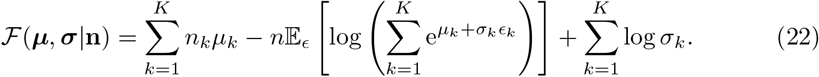

For given experimental counts **n**, maximizing this ELBO with respect to ***µ*** and ***σ*** yields estimates for the parameters of the middle `Bayesian` model in Fig 1.

Compared to the `Bayesian` model on its left, the current variational posterior requires twice as many parameters, which means that 2*K* parameters must be determined from only *K* observed counts. Because estimating more parameters than observations may appear problematic, we now analyze how this is achieved by the optimization of the ELBO.

#### 2.4.1 ELBO gradients

Setting the partial derivatives of the ELBO (22) with respect to *µ*_*k*_ and *σ*_*k*_ to zero, we find the following *variational-inference* (VI) conditions that need to hold at the maximum:

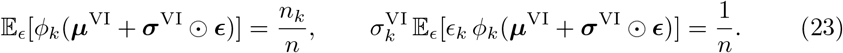

Note that the *k*th component of the *softmax* function,

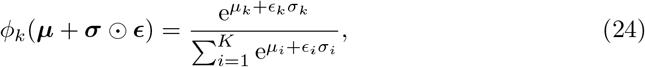

is a random variable due to the presence of ***ϵ*** in its argument. The ELBO gradients, however, are deterministic, since they involve expectations over ***ϵ***.

The first equality in (23) is directly analogous to the maximum-likelihood condition (15), requiring that the variant fraction predicted by the exponential model of growth (on the left) equals the empirical fraction observed in the data (on the right). The difference here is that the log abundances **y** are random variables described by a probability distribution, so this requirement applies to the expectation of the predicted fraction over this distribution.

Now consider scaling the counts of all variants by a common factor, for example due to an increase in sequencing depth. This scaling cancels in the right-hand side of the first equality in (23). The left-hand side must therefore remain invariant as well, implying that this equality carries no information about the absolute magnitude of the counts.

By contrast, the right-hand side of the second equality in (23) scales inversely with the total number of reads. Increasing all counts by a common factor thus forces the left-hand side to shrink proportionally. This in turn drives the standard deviations to decrease, though not necessarily by the same factor, since the expectation over ***ϵ*** also depends on ***σ***^VI^. In this way, the absolute sequencing depth is encoded into the variational standard deviations. Consequently, the *K* parameters ***σ*** play a role analogous to the single precision parameter *α* of the Dirichlet posterior of Sec 2.2.

A related observation is that in (23) the means ***µ*** always appear within the argument of the *softmax* function. As a result, their values can be shifted globally without affecting these optimality conditions. By contrast, the standard deviations ***σ*** also occur outside the *softmax* function, which prevents such global shifts and reflects their role in encoding the absolute scale of the counts. Accordingly, when maximizing the ELBO numerically, a gauge must be imposed on the means but not on the standard deviations.

This analysis clarifies the mechanism by which maximizing the ELBO extracts 2*K* variational parameters from *K* observations: the means encode relative proportions of the counts, while the standard deviations capture information about their absolute magnitudes. Because the expectations over the standard normal variables cannot be computed in closed form, however, the optimality conditions (23) cannot be solved analytically for the variational parameters. In Sec 2.4.2, we replace the expectation term in (22) with an analytical upper bound, thereby obtaining a tractable lower bound to the ELBO that can be maximized in closed form.

#### 2.4.2 Analytical lower bound to the ELBO

The Jensen inequality—stating that the expectation of a convex (concave) function is greater (smaller) than the function of the expectation [21]—provides a useful tool for deriving analytical bounds in variational inference. Applying it to the logarithm inside the expectation operator in (22) yields

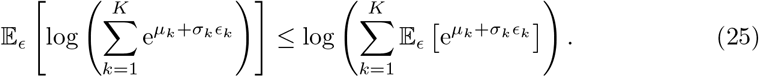

Bringing the expectation inside the logarithm allows us to calculate the expectations over the independent standard normal variables analytically:

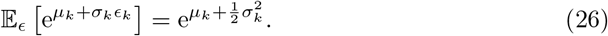

Replacing the second term of the ELBO (22) with its upper bound in (25) yields

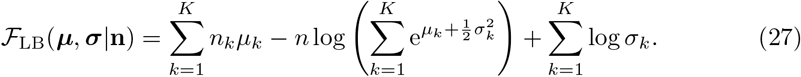

This expression provides a lower bound to the ELBO, since ℱ ≥ ℱ_LB_. It thus offers an analytically tractable surrogate objective for inferring the variational parameters by maximizing ℱ_LB_, instead of ℱ. Although this bound is not a controlled approximation to the ELBO, it has the advantage that its maximum can be obtained in closed form. Setting the gradients of *F*_LB_ to zero we find (see SI, Sec S3.3.1)

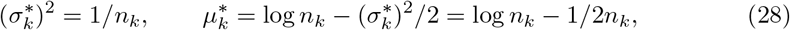

Where ∗ denotes the values of the parameter at the optimum of ℱ_LB_. The posterior variance of each variant is thus equal to the reciprocal of its counts.

Compared to the maximum-likelihood estimate 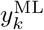 in (16), the mean 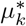in (28) contains a new additive term. Because we model *y*_*k*_ as being normally distributed, the abundance 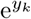 is log-normal. Hence, from (26), 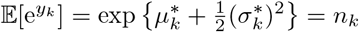. The new term therefore ensures that the expected abundance equals the observed counts.

To compare the inferred lower-bound posterior with the Dirichlet posterior of the variant fractions, we now use the equality (18). The means and variances of its left-hand side for the Dirichlet posterior are given in (10). The means and variances of its right-hand side for the lower-bound (LB) posterior are

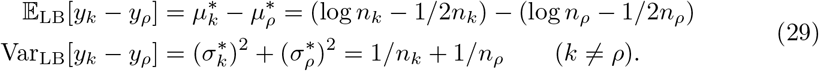

(The variance of the difference equals the sum of the individual variances due to the mean-field assumption of independence among the log abundances.) Since there are no any pseudo-counts in (29), we compare these to the Dirichlet expressions in (10) with 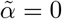. The two variances agree if the asymptotic expansion of *ψ*_1_(·) in (11) is terminated after the first term. The expectations, on the other hand, agree up to the second term in the expansion of *ψ*_0_(·). Thus, when the initial terms in the asymptotic expansions are sufficient, minimizing the analytical lower bound to the ELBO essentially recapitulates the Dirichlet analysis with zero pseudo-counts.

#### 2.4.3 Monte Carlo calculation of the ELBO

The above lower bound to the ELBO provides analytical insight, but its deviation from the ELBO cannot be reduced in a controlled way. Here we outline a numerical procedure for approximating the ELBO through Monte Carlo sampling.

From the perspective of sampling, the expectation term in the ELBO corresponds to the average over infinitely many independent random realizations. Specifically, if ***ϵ***^[*m*]^ denotes one random realization of the standard normal variables ***ϵ***, then

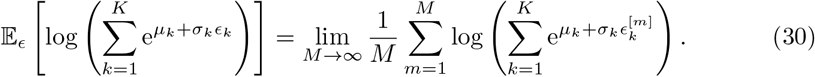

Thus, for a finite *M*, the right-hand side approximates the exact expectation.

Since, in general, the sample mean over another batch of *M* random realizations will be different, the approximate value will change stochastically every time it is calculated. Replacing the expectation term in the ELBO by the sample mean will thus lead to a stochastic approximation to the ELBO, which will continue to fluctuate even at the optimum. (This stochasticity complicates the assessment of convergence.) However, the approximation can be made increasingly more accurate by increasing the number *M* of stochastic samples, thus reducing the magnitude of the fluctuations at the cost of increased computational time.

When maximizing the ELBO numerically through gradient ascent, initializing the variational parameters close to their optimal values may greatly reduce the number of iteration steps. We therefore initialize the variational means and variances with the analytical values (28) from the Jensen lower bound.

#### 2.4.4 Numerical example

We illustrate the ideas discussed so far on a simple example with *K* = 4 variants. The starting fractions of the variants are chosen arbitrarily as

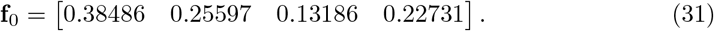

(These will change with time in Sec 2.5.2, where we introduce variant growth.)

To mimic experimental sequencing with 100*×* coverage, we pick the number of total read counts from a Poisson distribution with mean 400. Then, we use a multinomial distribution to partition the total counts according to the fractions (31). One realization of the counts is

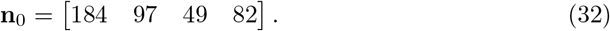

We now use these counts to estimate the variational parameters of the posterior Gaussian distributions.

Before reporting the estimated numerical values, it is necessary to decide on the gauge for the log abundances. We select the total initial abundance of all variants as a unit for measuring abundances, i.e., *N* (0) = 1, which means that the exponential sum of the means equals one, i.e.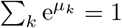.

The lower-bound parameters calculated from (28) are reported in the first row of Table 1. In the second row, we show the values obtained by maximizing the ELBO using *M* = 10^6^ random realizations of ***ϵ***. This relatively large sample size ensures an accurate approximation. Indeed, identical numerical values—within the three displayed decimal places—are obtained when the optimization is repeated with a different random seed.

**Table 1.** Variational estimates of initial log abundances and their uncertainties. True values are calculated from the fractions (31). Means and true values obey gauge condition 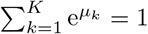.

| | $\mu$ | $\sigma$ |
| --- | --- | --- |
| Lower bound | $[-0.804 \quad -1.447 \quad -2.135 \quad -1.616]$ | $[0.074 \quad 0.102 \quad 0.143 \quad 0.110]$ |
| Monte Carlo ($M = 10^6$) | $[-0.804 \quad -1.447 \quad -2.135 \quad -1.616]$ | $[0.099 \quad 0.116 \quad 0.152 \quad 0.124]$ |
| True values | $[-0.955 \quad -1.363 \quad -2.026 \quad -1.481]$ | - |

To the reported precision, the means obtained by maximizing the ELBO coincide with those from the Jensen lower bound to the ELBO, However, the standard deviations from the lower bound are systematically smaller, thus underestimating the uncertainties compared to the maximization of the ELBO.

For comparison, the last row of Table 1 contains the true values of the initial log abundances calculated from (31), and shifted to the same gauge as the means. Since the variational estimates are based on a single realization of the counts, without any averaging over the stochasticity of the counting noise, close agreement between the estimated means and the true values is not expected. The discrepancies, however, should be captured by the estimated standard deviations ***σ***. Indeed, the true values fall within one standard deviation of the estimated means for all but the first variant, which lies within two standard deviations.

In summary, the analytical and numerical results presented here, demonstrate that variational inference offers a practical alternative to the Bayesian analysis with a Dirichlet posterior, despite the need to infer twice as many parameters from the data. In fact, having *K* precision parameters, instead of only one, could provide more flexibility for the purposes of uncertainty estimation.

### 2.5 Growth dynamics in terms of log abundances

In a growth competition experiment, the variant fractions change with time, and so the parameters of the multinomial probability distribution have different values at different sequencing time points. These values, however, are connected through the underlying growth dynamics of the variants. When a realistic model of the dynamics is available, the fractions at different time points can be expressed in terms of the model parameters. Consequently, estimates of these growth parameters can be obtained by jointly analyzing the counts across all time points.

In Sec 3, we critically analyze some of the existing approaches for achieving such joint analysis of the data through a model of variant growth. Here, we introduce three specific growth models—exponential, logistic and Gompertz [20]—for which we show how such analysis plays out.

For these three growth models, the variant cell numbers obey the differential equations

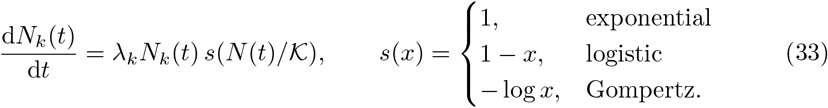

The specific growth rates *λ*_*k*_ are constants characterizing growth at the given experimental condition, and the function *s*(·) reflects the degree of saturation. When this saturation function equals zero, the cell numbers stop changing. For logistic and Gompertz growth, this happens when the total number of cells equals the carrying capacity 𝒦 (i.e., *x* = 1). Away from saturation (*x* ≪1), the saturation factor is less than one for logistic growth, but can be arbitrarily large for Gompertz growth, thus making the instantaneous growth rate in (33) much larger than the numerical value of the parameter *λ*_*k*_.

From the differential equations of the cell numbers, we obtain similar equations for the log abundances defined in (12):

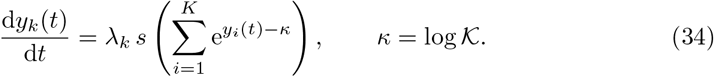

Integration from the initial time to the current sequencing time yields

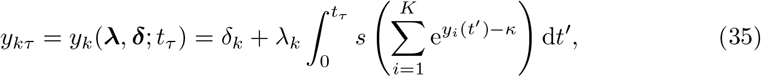

where we introduced the initial log abundances of the variants

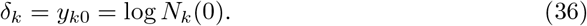

The growth dynamics of the *K* variants is therefore determined by the 2*K growth parameters* ***λ*** = (*λ*_1_, …, *λ*_*K*_) and **δ** = (δ_1_, …, δ_*K*_), assuming that the carrying capacity 𝒦 is known. Ultimately, the aim of the statistical analysis is to estimate all *K* growth rates from the numbers of sequencing reads over time, and to quantify the uncertainty of these estimates.

#### 2.5.1 Exponential growth

In the case of exponential growth, the saturation factor equals one, and the integral in (35) evaluates to *t*_*τ*_ . The log abundances therefore have the simple linear dependence

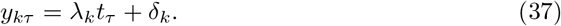

Substituting these into the *softmax* function (13) we find

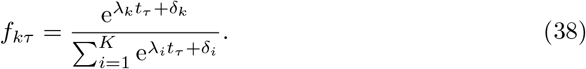

Inspection of this last result shows that both ***λ*** and **δ** can now be shifted independently without altering the variant fractions. Consequently, each set of parameters must be constrained separately in numerical work. For the initial log abundances, we use the gauge 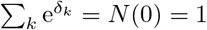, meaning that abundances are measured in units of the total initial population size. For the growth rates, we select the zero-sum gauge ∑_*k*_ *λ*_*k*_ =0.

A different gauge arises if the growth rate and initial log abundance of a reference variant are required to be zero. Rewriting the right-hand side of (18) for exponential growth yields

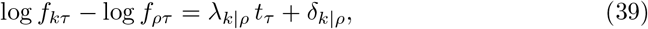

where we introduced the *relative* properties (i.e., with respect to the reference)

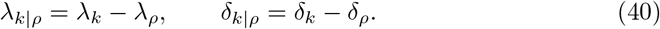

This choice of gauge, however, should not make the reference variant somehow special. In other words, the ultimate estimates of the growth parameters and their associated uncertainties should not depend on the identity of the reference variant.

#### 2.5.2 Numerical example

We now equip the illustrative example of four variants from Sec 2.4.4 with dynamics.

While the log abundances are available in closed form for exponential growth, they need to be integrated numerically for logistic and Gompertz growth, where analytical solutions exist only for a single variant (*K* = 1). The dynamics of the four variants under these three growth models, starting from the initial fractions (31), are shown in the top row of Fig 2.

**Fig 2.**
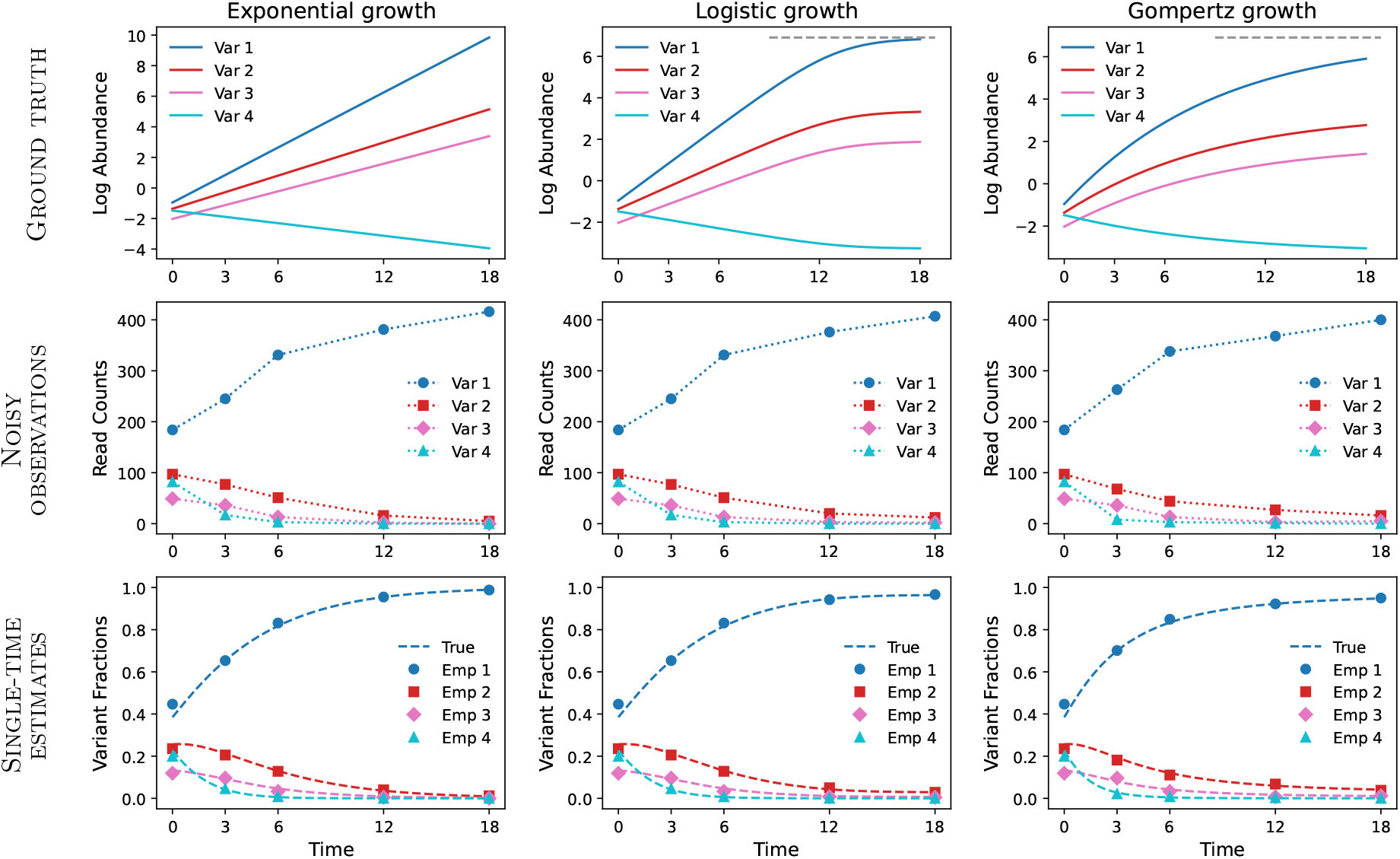
Illustrative example of exponential, logistic and Gompertz growth for *K* = 4 variants. Ground Truth: Synthetic growth trajectories of the variant log abundances. Exponential growth lines are calculated analytically using (37). Logistic and Gompertz growth are integrated numerically using Euler’s method with integration time step ten times smaller than the unit of time. Carrying capacity is indicated by horizontal dashed lines. Noisy Observations: Random multinomial counts generated according to the true fractions of the variants at the five specified time points. Symbols correspond to the numerical values in (43). Connecting dotted lines are for visual purposes only. Single-Time Estimates: Comparison between the empirical variant fractions in (6) calculated from the counts data (symbols, Emp), and the true fractions from which the counts were generated (dashed lines, True).

Following our choice of gauge, we set the initial total cell density to one, i.e., *N* (0) = 1. Dividing this density among the four variants according to the initial fractions (31) yields the initial log abundances (*cf* last row of Table 1)

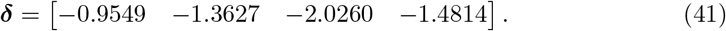

These are the *y*-intercepts of the Ground truth trajectories in Fig 2. The variant growth rates are arbitrarily chosen as

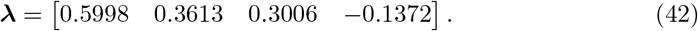

They correspond to the slopes of the lines in exponential growth, and have units of inverse time. (While the time unit here is arbitrary, it is intended to correspond to an hour.)

For the two models with saturation, we set the carrying capacity to be one thousand times larger than the initial cell density, i.e., 𝒦= 10^3^ and *κ* = 6.9. This value of *κ* is indicated by horizontal dashed lines in the top row of Fig 2. In the case of Gompertz growth, the saturation factor at the starting time evaluates to 6.9, which implies substantially higher initial growth rates compared to logistic growth, where the initial saturation factor is essentially 1. To ensure that Gompertz dynamics approach saturation on a similar timescale as logistic growth, we (arbitrarily) reduced ***λ*** in (42) by a factor of five for this growth model.

At the end of the displayed interval of 18 time units, the saturation factor equals 0.05 for logistic growth and 0.96 for Gompertz growth, showing that complete saturation (*s* = 0) is reached more closely by the logistic model. This is reflected in the maximum log abundances observed in the upper panels of Fig 2. Because exponential growth is unconstrained by carrying capacity, its log abundances are not bounded by 6.9 and can exceed this value.

The trajectories of logistic and Gompertz growth are obtained by integrating the differential equations (34) using the Euler method [30] with integration time step that is ten times smaller than the time unit, resulting in 180 steps between the initial and final times that are 18 time units apart. Decreasing the number of integration time steps would decrease the computational time.

Next, from the values of the Ground Truth log abundances in Fig 2, we compute variant fractions at the five specified time points (not spaced equally) and use these to generate random multinomial counts. As in Sec 2.4.4, the total counts at each time point are drawn from a Poisson distribution with mean 400, yielding roughly 100 counts per variant on average. For the three growth models, one realization of such multinomial counts is

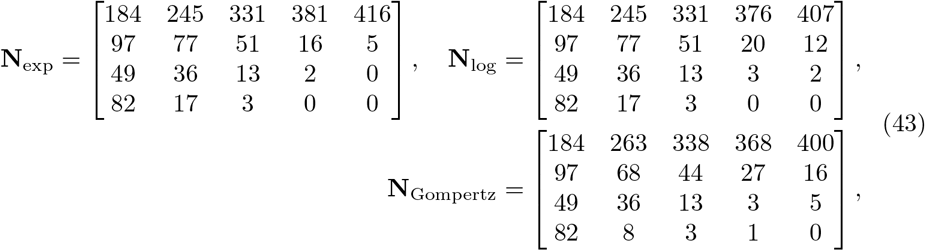

where rows correspond to variants and columns to time points. These values are visualized in the Noisy Observations row of Fig 2.

To be able to compare across the three models, we used the same random seed when generating the read counts. Consequently, the counts at the initial time point—first columns of the matrices in (43)—are identical across the three models. The counts at the subsequent time points depend both on the variant fractions determined by the Ground truth dynamics in Fig 2 and on the random draws from the multinomial distribution. Because logistic growth is practically indistinguishable from exponential growth away from saturation, the two models yield nearly identical variant fractions at early times. In our example, their counts remain identical at the second and third time points, a consequence of having used the same random seeds in generating the multinomial samples from the variant fractions. By contrast, the counts for Gompertz growth begin to deviate immediately after the first time point (except for the third variant).

From the synthetic counts (43), we compute the empirical variant fractions *n*_*kτ*_ */n*_*τ*_, which are the maximum-likelihood estimates (6). These are plotted as symbols in the bottom row of Fig 2. Not surprisingly, they closely track the true generating fractions (dashed lines). Note that these maximum-likelihood estimates do not incorporate any temporal information from the growth model. (This is achieved below in Sec 3.2.)

In this example, the fastest-growing variant rapidly dominates the population, eventually accounting for nearly all observed counts. As a result, the counts across variants begin to differ by several orders of magnitude, with some variants disappearing altogether. Such pronounced imbalances are also expected in large-scale growth competition experiments involving thousands of variants.

Because rare variants are particularly susceptible to counting noise, a principled approach to estimate model parameters together with their associated uncertainties is needed. In Sec 3.3, we employ Bayesian inference as a general framework for parameter estimation and uncertainty quantification [11]. Before that, in Sec 3.1, we use Bayesian analysis to derive variance-based weight factors for the purposes of least-squares fitting.

## 3 Results

In this section, we address the question of how to best integrate the temporal dimension of the sequencing counts data through a model of growth. We start by revisiting the least-squares fitting strategy (Sec 3.1), used for example in Enrich2 [9]. Then we move on to maximum-likelihood estimation (Sec 3.2) and variational inference (Sec 3.3) [11]. While these three subsections focus on exponential growth, in Sec 3.4 we extend the maximum-likelihood and variational-inference approaches to growth models with saturation.

### 3.1 Weighted least-squares fitting

The equalities (38) and (39) constitute a natural starting point for fitting the parameters of the analytical growth model (on their right-hand sides) to the experimental sequencing data. To this end, their left-hand sides are first estimated from the read counts at each sequencing time point, and then the cumulative error between the estimates across time points and the predictions of the growth model is minimized.

This approach implicitly assumes that the errors at each time point are normally distributed and have identical variances. The condition of constant variance can be relaxed by normalizing the squared errors at different times by their respective variances, leading to a *weighted* least-squares fitting, which requires estimating both the means and variances of the left-hand sides of (38) and (39) from the data. For the Dirichlet posterior of Sec 2.2, the needed means and variances are given in (9) and (10).

#### 3.1.1 Fitting the logarithms of the fractions

Starting with (39), we construct the weighted mean-squared error

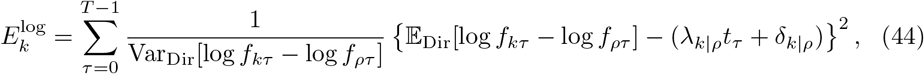

where the squared deviations between the posterior means and the growth model are normalized by the posterior variances. This weighting accounts for heterogeneity in statistical uncertainty across time points, ensuring that more precise estimates contribute more strongly to parameter inference [9].

Substituting the means and variances from (10), and using a single pseudo-count parameter 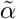, we rewrite (44) as

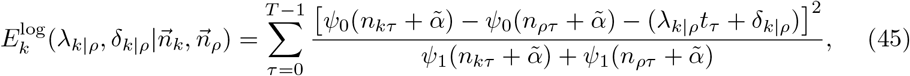

where 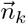 and 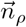 denote the counts of variants *k* and *ρ* across all sequencing time points. To the best of our knowledge, this expression has not been used in the literature.

What has been used, for example in Enrich2 [9], is its large-count limit. Retaining only the first terms in the expansions (11), the error (45) can be approximated as

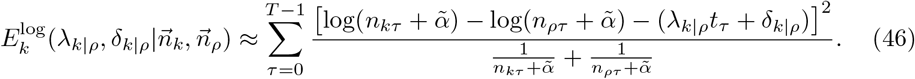

For the choice 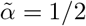, this becomes the Enrich2 error [9].

The error (45) is for variant *k*, and depends only on the relative growth parameters *λ*_*k*|*ρ*_ and δ_*k*|*ρ*_ defined in (40). To estimate all growth parameters, a separate minimization should be preformed for each non-reference variant. Alternatively, the individual variant errors can be added together, and the total error can be minimized simultaneously with respect to all growth parameters, keeping those of the reference variant equal to zero.

For the illustrative example with four variants (Sec 2.5.2), we minimized the sum of the errors (45), with variant *ρ* = 2 as reference and 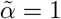 (i.e., one additional pseudo-count). The fits for the three non-reference variants are shown in the upper row of Fig 3A (red lines). In addition, we show the target posterior means and standard deviations (orange points and error bars). When the standard deviation is large, the squared difference between the linear model and the posterior mean contributes less to the total error. As expected, the largest standard deviations correspond to zero variant counts (last point of variant 3 and last two points of variant 4). In these cases, the magnitudes of the standard deviations are mainly dictated by the pseudo-count parameter 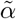.

**Fig 3.**
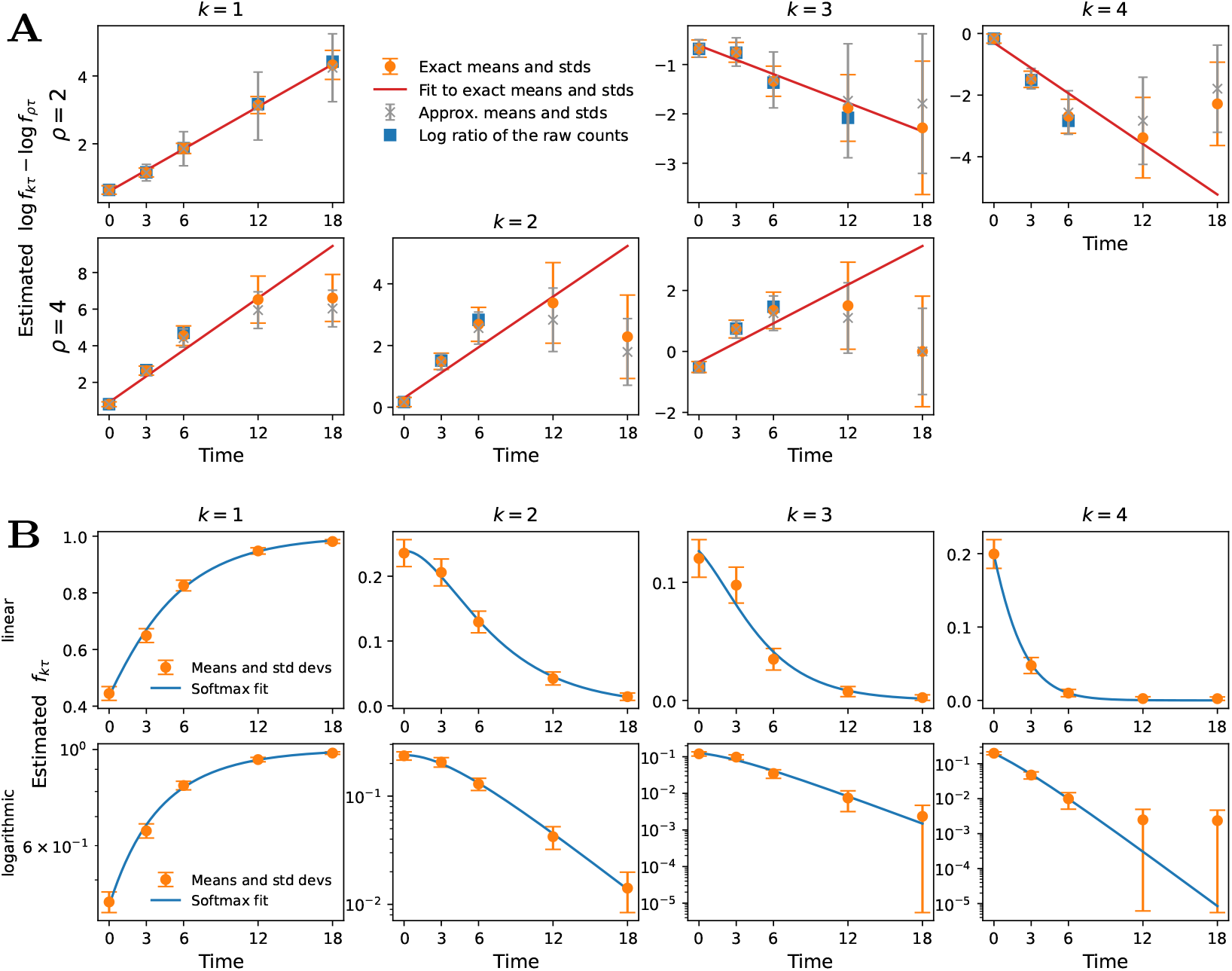
Weighted least-squares fits for exponential growth. **A**: Linear fits with variant 2 (top row) or variant 4 (bottom row) used as reference. Exact means and standard deviations from (10) are in orange and approximations used in the error (46) are in gray. Blue squares show log *n*_*kτ*_ ∗log *n*_*ρτ*_, which are the maximum-likelihood estimates without any pseudo-counts. **B**: *Softmax* fits with variant fractions displayed on linear (top row) or logarithmic (bottom row) vertical scales. Although variant 2 is used as a reference, identical fits are obtained when *ρ* is any other variant.

For comparison, in Fig 3A we also show the approximate means and standard deviations (gray crosses and error bars) used in the error (46), again calculated with a uniform prior 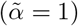. To assess the effect of the pseudo-counts, we additionally plot the maximum-likelihood estimates log 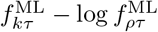 (blue squares), which do not include any pseudo-counts. Note that these are available only when the counts of both the analyzed variant and the reference variant are non-zero, since otherwise the logarithms evaluate to minus infinity. For the non-zero counts, both the approximate means (gray crosses) and the log ratios of the raw counts (blue squares) agree well with the exact posterior means (orange points). Not surprisingly, the discrepancy between the exact and approximate posterior means is larger for zero counts, since the approximation obtained by terminating the asymptotic expansions at the first term applies for large numbers.

Focusing now on the proximity of the fitted line to the target points, from the top-right panel in Fig 3A we observe that the last two points of variant 4, which lack any actual counts, pull the fitted line away from the linear trend of the first three points. As a result, at three out of the five points—including two points with non-zero counts—the fitted line is more than one standard deviation away from the means.

The fit of the most abundant variant 1, shown in the top-left panel of Fig 3A, is perfect. What is interesting in this case is the increase of the standard deviations at the last two time points, in spite of the fact that the counts of variant 1 increase steadily with time and attain their largest values there. Somehow counterintuitively, the largest counts of variant 1 at the last two time points contribute to the total error with smaller weights than the comparatively smaller counts at the first three time points. The explanation of this seemingly strange phenomenon lies in the fact that the error bars correspond to the variances in (10), which involve the reference variant.

To better demonstrate the influence of the reference variant on the fits, in the second row of Fig 3A we show the same analysis but using *ρ* = 4 as a reference. In this case, even for the most abundant variant 1 the fitted line lies more than one standard deviation from two of the five target points shown in orange. For all variants, the last two points, where the reference variant has zero counts, visibly deviate from the linear trend of the first three points.

In the first two rows of Table 2, we give the numerical values of the growth rates obtained from the linear fits in Fig 3A. Column ***λ*** shows the values shifted globally to a zero-sum gauge, and column Δ*λ* gives the differences between two successive variants.

**Table 2.** Growth rates from weighted least-squares fits to exponential growth. Errors *E*^log^ (45) and *E*^frac^ (48) are minimized with variant *ρ* used as reference. Entries of Δ*λ* are Δ*λ*_*k*_ = *λ*_*k*_ ∗ *λ*_*k*+1_, with *k* = 1, …, *K* ∗ 1.

| | $\lambda$ (zero-sum gauge) | | | | $\Delta\lambda$ | | | |
| --- | --- | --- | --- | --- | --- | --- | --- | --- |
| $E^{\log}(\rho = 2)$ | 0.249 | 0.041 | -0.056 | -0.233 | 0.208 | 0.097 | 0.177 | Fig 3A, top |
| $E^{\log}(\rho = 4)$ | 0.235 | 0.034 | -0.028 | -0.240 | 0.201 | 0.062 | 0.211 | Fig 3A, bottom |
| $E^{\text{frac}}(\rho = 2)$ | 0.275 | 0.071 | -0.017 | -0.329 | 0.204 | 0.088 | 0.312 | Fig 3B |
| $E^{\text{frac}}(\rho = 4)$ | 0.275 | 0.071 | -0.017 | -0.329 | 0.204 | 0.088 | 0.312 | |

The dramatic effect of the reference variant on the fits, as observed in this simple example, contradicts our expectation that the selection of reference is needed only to fix the trivial global shift in growth rates and initial log abundances.

#### 3.1.2 Fitting the fractions

A different weighted mean-squared error can be formed from the *softmax* identity (38):

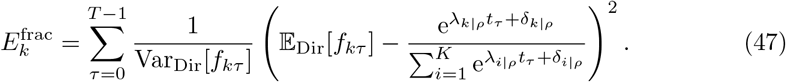

For consistency with the error (44), we have fixed the gauge of the *softmax* function in (47) by subtracting the growth rate and initial log abundance of a reference variant, thus again working with values relative to *ρ*.

Differently from before, the error of variant *k* in (47) depends on the growth parameters of all variants, since these are used in the denominator of the *softmax* function for the purposes of normalization. To minimize all parameters simultaneously, we add the individual errors, giving equal weights to all variants. After substituting the means and variances from (9), this total error becomes

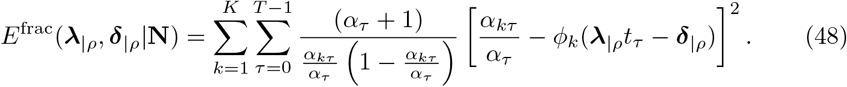

The dependence on the raw counts **N** comes from the posterior Dirichlet parameters *α*_*kτ*_ according to (8). This error also appears to be new.

For the example with four variants, we minimized (48) using 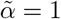, again selecting variant *ρ* = 2 as a reference. The result is shown in Fig 3B. On the vertical axes we now plot the variant fractions, since their means (orange points) are the target values of the fit. The same fractions are plotted on a linear (top) and logarithmic (bottom) scale. The fitted analytical expression (blue curves) is no longer linear but is given by the *softmax* function.

Looking at the top row of Fig 3B first, we see that the error bars of variant 1 decrease with time, reflecting the increase of its counts. Interestingly, the error bars of all other variants also decrease with time, although their counts are gradually depleted, and eventually become zero for variants 3 and 4. As a result, the zero counts of these two variants have tiny error bars, and thus contribute to the total error with larger weights, in contrast to the linear fits in the top row of Fig 3A.

However, it is not fair to directly compare the error bars in the upper rows of Figs 3A and 3B, because the latter shows the variant fractions and the former the logarithms of these fractions. To be able to visually compare the two alternative fitting strategies, in the lower half of Fig 3B we show the same variant fractions on a logarithmic scale. These lower panels are therefore directly comparable to those in Fig 3A, where the logarithms of the fractions are on a linear scale.

Satisfyingly, for variant 4 in the lower-right panel of Fig 3B, the error bars of the last two time points are much larger than the error bars of the first three points, in qualitative agreement with the upper-right panel of Fig 3A. However, while in Fig 3A these zero-count points were pulling the fitted red line away from the trend of the first three points, here the fitted blue curve not only goes straight through the first three points, but also remains within one standard deviation of the last two points.

In the case of variant 1 in the lower-left panel of Fig 3B, the error bars at later times are smaller than those at earlier times, which is the opposite of the upper-left panel of Fig 3A for the linear fit of this variant. Recall, however, that the error bars in the linear fit depend also on the counts of the reference variant, while those of the *softmax* fit do not.

This comparison shows that the errors (45) and (48) treat the reference variant in fundamentally different ways. While the linear fits depend strongly on the choice of reference, as demonstrated in Fig 3A, the *softmax* fits do not. Indeed, identical fits are obtained when the minimization is repeated with *ρ* = 4, even though its last two counts are zero. (The fitted growth rates are given in the last two rows of Table 2.) For the purposes of fitting exponential growth, therefore, the *softmax* fit appears to be more satisfying than the linear fit.

The dramatically different performance of the two alternatives stems from the different normalization choices. The *softmax* transformation (38), on which the error (47) is based, cancels the arbitrary global shift of the growth parameters through the division by the sum over all variants. Correspondingly, the estimates of the fractions contain normalization by *α*_*τ*_, which also sums over all variants. The motivation for arriving at (39), on which the linear fit is based, was to cancel the normalization by all variants. Subtracting the reference for that purpose, amounts to normalization by the reference variant, which becomes evident when the difference of logarithms is written as log(*f*_*kτ*_ */f*_*ρτ*_). Analytically, these two normalizations are as good as any other way of fixing the gauge. From the perspective of estimation, however, normalization by a single variant appears to be more susceptible to counting noise than normalization by all variants.

Historically, a linear fit may have been preferred over a non-linear *softmax* fit for computational reasons, since minimizing the error in linear regression requires a single matrix inversion. By contrast, minimizing the *softmax* error requires iterative numerical optimization, such as gradient descent. Moreover, the linear fit can be performed independently for each variant, optimizing only two parameters at a time, whereas the *softmax* fit requires the simultaneous optimization of all 2*K* growth parameters. With the advent of modern computational resources and widespread availability of automatic differentiation libraries [19], these computational disadvantages of the weighted *softmax* regression are no longer prohibitive—even when analyzing thousands of variants.

### 3.2 Maximum-likelihood estimation of the growth parameters

The weighted least-squares approach to estimating growth parameters—including its *softmax* variant—suffers from two main shortcomings. First, although Bayesian reasoning is invoked to construct the weight factors, the method ultimately produces only point estimates of the growth rates, with no quantitative measure of their uncertainty. This limitation arises because Bayesian inference is applied too early in the analysis pipeline: instead of capturing uncertainty in the final growth-rate estimates, it is directed toward the intermediate variant fractions. Second, the approach aggregates independent maximum-likelihood estimates from individual time points by minimizing a mean-squared error, thereby treating the temporal structure of the data only indirectly, as an afterthought. Yet the time axis can be incorporated directly within the maximum-likelihood framework itself, as we demonstrate in this section.

The Bayesian treatment of the growth parameters in Sec 3.3 builds on the all-time likelihood function that we introduce here.

#### 3.2.1 Combining all time points

To obtain maximum-likelihood estimates of the growth parameters directly, the later must appear explicitly in the likelihood of the probabilistic model. However, the multinomial distribution that we use to represent counting noise is parametrized in terms of variant fractions. We therefore reparametrize this distribution in two successive steps.

First, we assume that the counting noise at one sequencing time is independent of that at any other time, which allows us to sum the single-time log likelihoods (14):

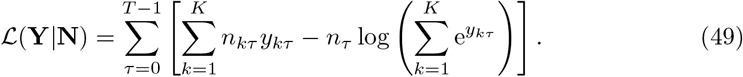

Summing across time, however, does not by itself achieve the desired integration. Indeed, maximizing (49) with respect to the log abundances *y*_*kτ*_ recovers the single-time maximum-likelihood conditions (15).

In the second step, we incorporate the growth model to express the time-dependent log abundances in terms of the constant growth parameters. This yields the All TIMES-`Likelihood` model in Fig 1, whose likelihood can now be maximized directly with respect to ***λ*** and **δ**.

In the case of exponential growth, the resulting log likelihood is

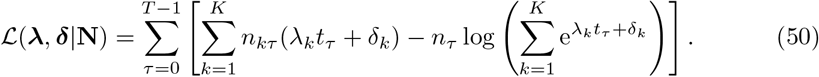

Maximizing (50) estimates 2*K* parameter from *K × T* observations, corresponding to *T/*2 observed counts per estimated parameter.

To understand how maximum-likelihood estimation integrates information across time, it is instructive to examine the gradients of (50) with respect to the growth parameters. Differentiating with respect to *λ*_*k*_ and δ_*k*_, and setting the derivatives to zero, we find that the optimal ***λ***^ML^ and **δ**^ML^ must satisfy

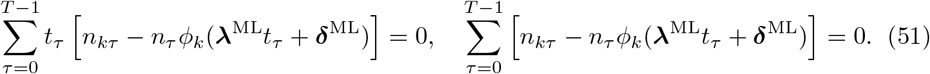

In both equations, the bracketed term compares the actual counts of a variant, *n*_*kτ*_, to its counts predicted on the basis of the growth model, *n*_*τ*_ *ϕ*_*k*_(***λ****t*_*τ*_ + **δ**). This term vanishes when

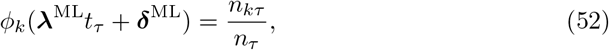

which is exactly the maximum-likelihood estimate (15) obtained by analyzing each time point independently. When the data are analyzed jointly across all time points, however, condition (52) need not hold at every *t*_*τ*_ . Instead, the requirement is that only the time-summed equalities in (51) are satisfied. In other words, joint estimation allows positive and negative deviations of the predicted and observed counts at individual time points to balance each other out across time.

The same type of balancing occurs also for the gradients of the mean-squared error in least-squares fitting. The key difference is that least-squares assumes Gaussian noise, with variances adjusted by explicitly assigning weights to the errors at different time points. By contrast, the conditions in (51) are derived directly from the multinomial sampling model, and the appropriate weighting emerges naturally rather than being imposed externally.

Occasionally, the counts of certain variants may drop to zero at specific time points, either because the variants go extinct or because sequencing depth is insufficient to capture very small fractions. This occurs, for example, with variants 3 and 4 in our illustrative data (43). In the Bayesian analysis of the previous section, zeros are avoided by adding a constant number of pseudo-counts to the observed data. However, the maximum-likelihood estimation procedure does not include pseudo-counts. We therefore examine the effect of zero counts on the conditions (51).

When *n*_*kτ*_ = 0, the corresponding term in the square brackets contributes negatively to both sums in (51), since the *softmax* output is strictly positive. To satisfy the equalities, these negative contributions must be compensated by positive contributions from other time points where the predicted counts fall below the observed counts. For such cancellations to occur, however, at least one nonzero count of the variant is required. In fact, because two parameters are estimated per variant, at least two nonzero time points are necessary. If fewer than two are available, zero counts cannot be analyzed without the introduction of pseudo-counts.

In our illustrative four-variant example, we maximized the all-time log likelihood (50) using gradient ascent. After each iteration step, the optimization parameters were adjusted to enforce the gauge conditions: growth rates were constrained to sum to zero, and the exponential sum of the initial log abundances was normalized to one, corresponding to *N* (0) = 1. The final growth rates are shown in the second row of Table 3 (ML, *T* = 5). (The first row repeats the weighted least-squares growth rates from the last two rows of Table 2.)

**Table 3.** Estimates of the variant growth rates and their uncertainties for exponential growth. Counts at all five time points are used for *T* = 5, while only those at *t*_0_ = 0 and *t*_2_ = 6 are used for *T* = 2. (WLSQ: weighted least-squares with *E*^frac^; ML: maximum likelihood; VI: variational inference with ELBO; LB: analytical lower bound to ELBO.)

| | $\lambda$ or $\mu^\lambda$ (zero-sum gauge) | $\sigma^\lambda$ (uncertainty) | |
| --- | --- | --- | --- |
| WLSQ ($T = 5$) | [0.275 0.071 -0.017 -0.329] | - | Fig 3B |
| ML ($T = 5$) | [0.297 0.084 -0.025 -0.356] | - | Fig 4, left |
| ML ($T = 2$) | [0.293 0.088 -0.026 -0.356] | - | Eq (54) |
| VI ($T = 5$) | [0.301 0.086 -0.026 -0.362] | [0.012 0.013 0.029 0.062] | Fig 5, left |
| LB ($T = 5$) | [0.300 0.087 -0.025 -0.362] | [0.002 0.012 0.028 0.061] | Fig 5, right |
| LB ($T = 2$) | [0.302 0.096 -0.023 -0.375] | [0.009 0.023 0.046 0.096] | Eq (64) |
| True $\lambda$ from (42) | [0.319 0.080 0.019 -0.418] | - | |

To visualize how well the estimated ***λ***^ML^ and **δ**^ML^ compare with the data, on the left side of Fig 4 we show the trajectories of the variant fractions calculated from these estimates. As expected, the estimated fractions (solid lines) deviate from the true ones (dashed lines) in order to better match the empirical fractions (symbols), which are the single-time maximum-likelihood estimates (6).

**Fig 4.**
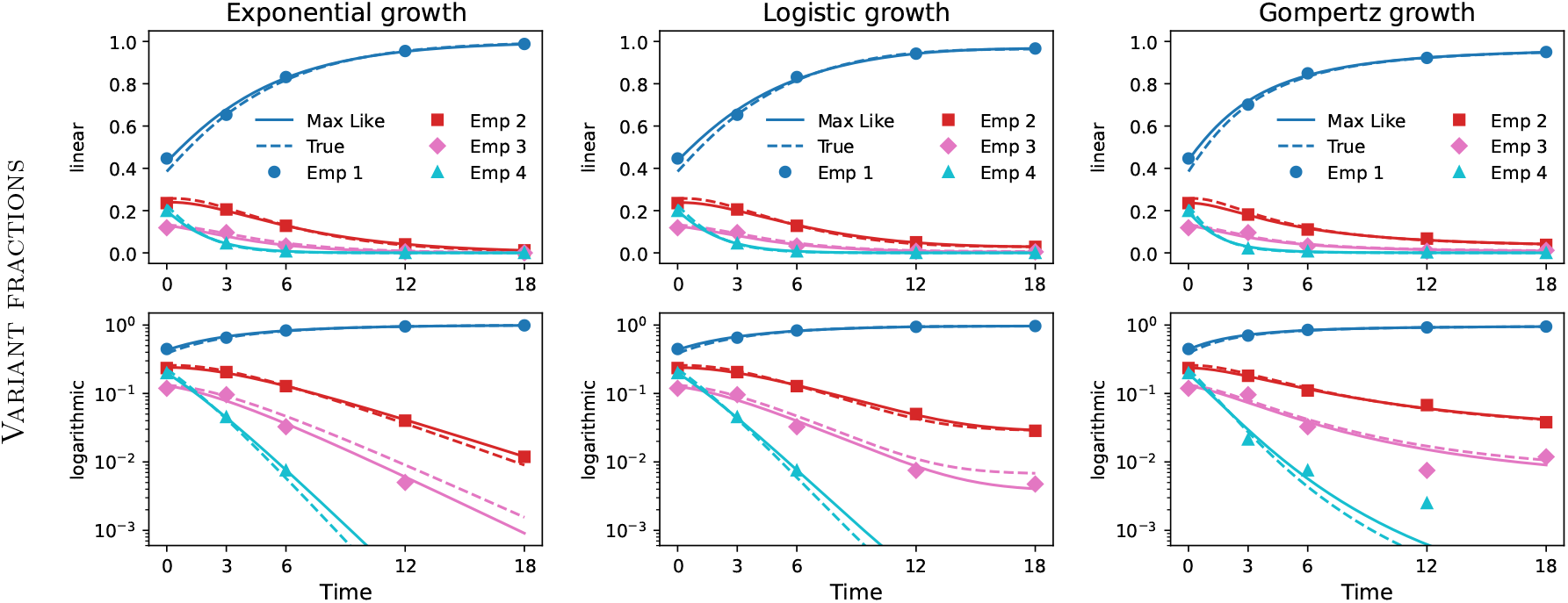
Time trajectories of variant fractions from maximum-likelihood estimates of the growth parameters. Calculations based on the all-times maximum-likelihood (ML Time) estimates of the growth rates and initial log abundances are compared to the true (dashed lines) and empirical (symbols) fractions. For each growth model, the fractions are plotted on a linear (top) and logarithmic (bottom) vertical scale to better resolve both many and few counts.

Although Gompertz growth is discussed in Sec 3.4, we draw attention at this point to the identical analysis presented on the right side of Fig 4. The curves for variants 3 (cyan) and 4 (pink) in the lower-right panel nicely illustrate that counts with lower abundance—or zeros—contribute less than higher counts. This intuitive weighting emerges automatically from the maximum-likelihood framework, in contrast to least-squares fitting, where weights had to be imposed externally.

#### 3.2.2 Two sequencing time points only

Given that many studies sequence only at the beginning and end of a competition experiment [5], we now analyze the zero-gradient conditions (51) for the case *T* = 2. Because of the multiplication by *t*_*τ*_ in the first equality, the initial time point (*t*_0_ = 0) does not contribute to this sum. Hence, the condition (52) must be satisfied entirely by the second time point. Once this holds, however, the second time point contributes nothing to the second equality in (51), since its bracketed term vanishes. Consequently, the first time point alone also satisfies (52). The case *T* = 2 therefore reduces to performing separate maximum-likelihood estimations at the two time points, without any integration of information across time.

Written explicitly, the separate conditions (52) at the first and second time points are

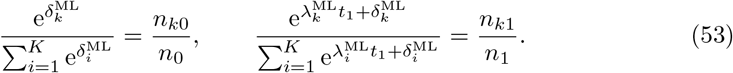

The first equality does not contain the growth rates because they are multiplied by *t*_0_ = 0. As a result, the initial log abundances are estimated only from the initial counts (up to a global shift). With **δ**^ML^ determined from the first time point, the equality at the second time point fixes the variant growth rates (up to another global shift). Thus, as noted earlier, for *T* = 2 the observed read counts are only just sufficient to determine the *y*-intercepts and the growth rates, leaving no redundancy for averaging over the counting noise. The only way to go beyond this bare minimum is to sequence at more than two points.

The equalities (53) can be solved for the maximum-likelihood estimates of the growth parameters. Since the latter are identifiable only up to arbitrary global shifts, we discard additive contributions that are common to all variants (i.e., terms without the index *k*), to arrive at

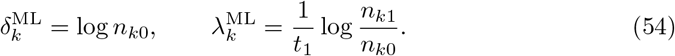

Interestingly, the estimator of the growth rate depends solely on the counts of variant *k* at *t*_0_ and *t*_1_, irrespective of whether the total counts at these two time points differ.

Once obtained, the estimated growth rates and initial log abundances can be shifted collectively to any other desired gauge.

As an example, we use (54) to calculate growth rates only from the counts at the first and third time points of our synthetic data (43). (The third time point is the last instance where the counts of all variants are non-zero.) The resulting numerical values are given in the third row of Table 3 (ML, *T* = 2). They are practically identical to the maximum-likelihood estimates based on the counts at all five time points, shown above them.

Using the estimators (54) and the definition of relative growth parameters (40), one can switch to a gauge where the parameters of a reference variant, *ρ*, are set to zero:

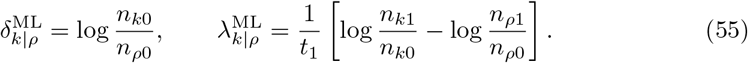

The expression for 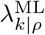 is the standard formula used to estimate relative growth rates from sequencing data at two time points. Our analysis demonstrates that this estimator yields the maximum-likelihood estimate of the relative growth rate for the case *T* = 2.

Our alternative growth-rate estimator in (54) involves only the counts of variant *k*, whereas the conventional estimator in (55) also depends on the counts of the reference variant. This distinction has direct implications for the statistical uncertainties of the two estimators, as we discuss in Sec 3.3.2.

#### 3.3 Variational inference for the growth parameters

Promoting the growth parameters of the previous subsection to random variables, and approximating the posterior once again by a product of independent normal distributions, yields the All Times–`Bayesian` model in Fig 1.

The variational parameters of this new model are inferred from the data by maximizing the ELBO

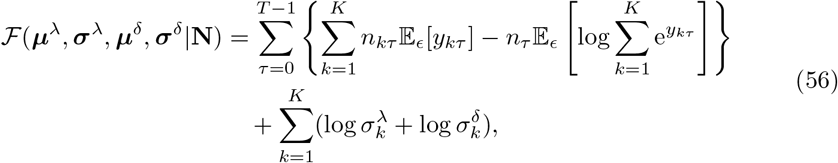

where the dependence on the means and standard deviations is through the log abundances *y*_*kτ*_, which also depend on the standard normal variables 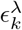 and 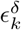 . The operator 𝔼_*ϵ*_[·] now takes expectation over both ***ϵ***^*λ*^ and ***ϵ***^δ^.

As in Sec 2.4, the final term of the ELBO (56) represents the entropy of the posterior, now receiving contributions from both the growth rates and the initial log abundances, as indicated by the superscripts. The first term is the “mean energy,” which now includes a sum over all time points. This ELBO does not explicitly incorporate the model of growth yet, and so can be combined with any growth dynamics, including the saturating growth that we analyze in Sec 3.4.

In the case of exponential growth, 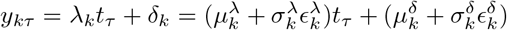 so the first expectation in (56) can be evaluated analytically:

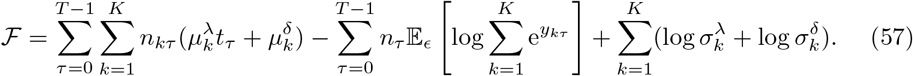

As a result, only the second term retains an expectation operator, mirroring the structure of the single-time ELBO (22).

As before, this expectation can be approximated stochastically by drawing *M* samples of (***ϵ***^*λ*^, ***ϵ***^δ^) and taking the empirical mean. Alternatively, it can be replaced with an analytical upper bound from Jensen’s inequality, thereby obtaining a fully analytical lower bound on the ELBO, which can be maximized instead. This second approach is pursued below in Sec 3.3.1, after first evaluating the ELBO numerically for our illustrative example with four variants.

We calculated the ELBO (57) by generating *M* = 10^4^ random realizations per iteration step, using automatic differentiation for the gradients. At the end of each step, the means ***µ***^*λ*^ were shifted such that they summed to zero and the means ***µ***^δ^ such that their exponential sum equaled one. To accelerate convergence toward the optimum, the variational means were initialized with the maximum-likelihood estimates ***λ***^ML^ and **δ**^ML^, while the standard deviations were initialized as

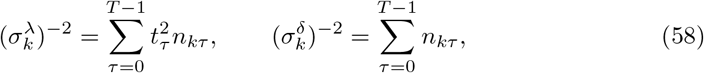

a choice motivated later in (62). Convergence was assessed visually by verifying that the ELBO stabilized into stochastic fluctuations around a steady baseline (see SI, Fig S4). Final estimates were obtained by averaging the fluctuating parameter values over the last 100 optimization steps.

The inferred growth-rate means and standard deviations are reported in the fourth row of Table 3 (VI, *T* = 5). On the left side of Fig 5, we show the posterior Gaussian distributions of the growth rates (top) and initial log abundances (bottom). We also show the maximum-likelihood estimates from Sec 3.2.1 (dashed colored lines) and the true parameter values used to generate the synthetic data (solid black lines).

**Fig 5.**
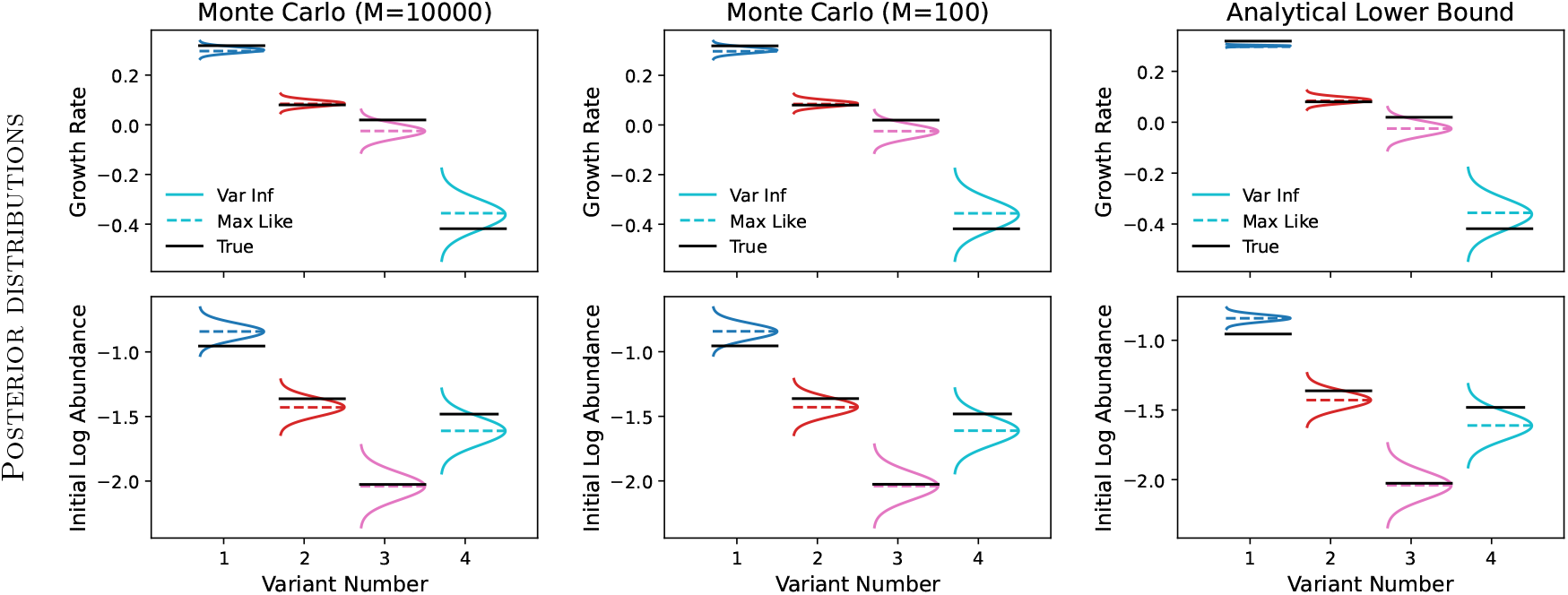
Variational posteriors from Monte Carlo and analytical approximation to ELBO under exponential growth. The Gaussians are shown in the interval *µ*_*k*_ *±*3*σ*_*k*_. True values of the growth parameters are indicated with black lines and maximum-likelihood estimates (Max Like) with dashed lines. Decreasing the number of random realizations in the stochastic approximation from 10^4^ to 10^2^ does not change the results. In comparison, minimizing the analytical lower bound to the ELBO underestimates the uncertainties, especially for the most abundant variant 1.

Visually, the uncertainties of the more abundant variants 1 and 2 are comparable and smaller than those of the less abundant variants 3 and 4, although the difference is more pronounced for the growth rates [Table 3 (VI, *T* = 5)] than for the initial log abundances. Additionally, the growth-rate uncertainty of variant 4, which has zero counts at the last two time points, is substantially larger than that of variant 3, which is zero only at the last time point.

Given that a sample size of *M* = 10^4^ may become prohibitively large for thousands of variants, we explored the possibility of reducing this number. The middle panels in Fig 5 demonstrate that approximating the expectation with as little as *M* = 100 random samples yields identical results. This smaller number is used in the subsequent numerical examples.

#### 3.3.1 Analytical lower bound to the ELBO

For further analytical insight, we now analyze the Jensen lower bound to the exponential-growth ELBO (57):

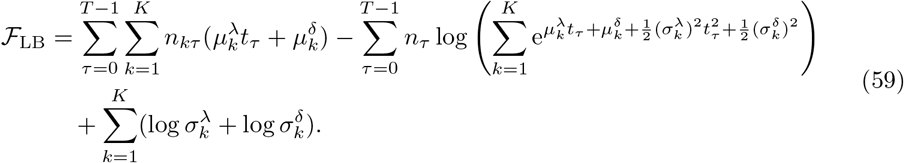

Its partial derivatives with respect to the means yield the optimality conditions

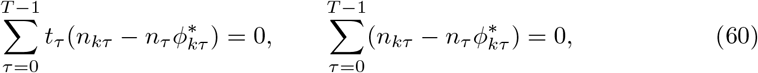

where

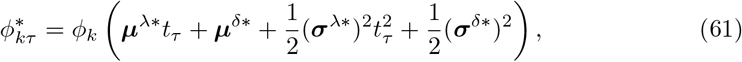

and denotes values at the optimum of ℱ_LB_. Although these gradient conditions take the same form as those in (51), derived from the all-time log likelihood, the argument of the *softmax* function here includes means and variances.

Further conditions are obtained from the partial derivatives of ℱ_LB_ with respect to the standard deviations:

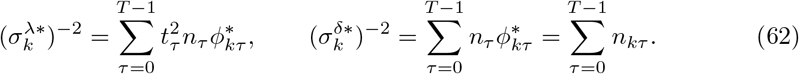

According to the last equality, the inverse variance (also known as *precision*) of the initial log abundance δ_*k*_ equals the sum of the counts of that variant across all time points. The precision of *λ*_*k*_ in (62) depends on 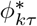, which is yet to be evaluated. Nonetheless, we recognize that 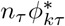 corresponds to the predicted counts of variant *k* at *t*_*τ*_ . Motivated by these results, when maximizing the ELBO numerically, we initialize the standard deviations as given in (58).

While the inverse proportionality between variance and number of counts is characteristic for counting processes like Poisson and multinomial, the precisions in (62) are directly for the growth parameters, and show how to aggregate the counts across time points. Admittedly, these analytical results are for the lower bound to the ELBO, and are expected to be quantitatively different from the posterior variances obtained by maximizing the ELBO itself.

To assess this difference for the illustrative example with four variants, we maximized ℱ_LB_ numerically using gradient ascent. The resulting posterior distributions are shown on the right of Fig 5; the corresponding growth-rate parameters are given in the fifth row of Table 3 (LB, *T* = 5). For the most-abundant variant 1, the standard deviations obtained by maximizing the lower bound are substantially narrower that those obtained by maximizing the ELBO. This underestimation of the statistical uncertainty is in line with the single-time result for *t*_0_ = 0 in Table 1. For the other three variants in Fig 5, however, the underestimation is visually undetectable. Indeed, the standard deviations of the last three variants in Table 3 (LB, *T* = 5) are essentially identical to the ones above them (VI, *T* = 5).

This numerical example demonstrates that the uncertainty of highly abundant variants may be significantly underestimated when inferred from the lower bound to the ELBO. Keeping this limitation in mind, we further pursue the maximization of ℱ_LB_ in for the special case of only two sequencing time points.

#### 3.3.2 Sequencing at only two time points

The optimality conditions (60) and (62) admit further simplification in the case of only two sequencing time points, analogously to the situation in Sec 3.2.2. Following the same logic as there, we find that for *T* = 2

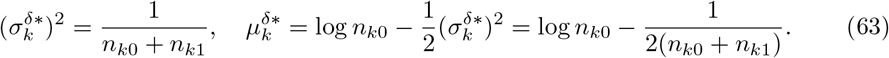

Compared to the maximum-likelihood estimate δ^ML^ in (54), the last term in (63) is new. It reflects the entropic correction to the mean due to the variance. Because we model δ_*k*_ as being normally distributed, the initial abundance 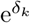 is log-normal. Hence, from (26), 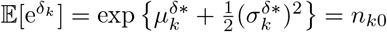, so the new term on the right-hand side of (63) ensures that the mean abundance equals the observed initial counts.

Due to the presence of the counts at the second time point *t*_1_, the mean in (63) is also different from the variational mean in (28) calculated at a single time point.

For the growth rate in the case of sequencing at only two time points, we find from (60) and (62) that

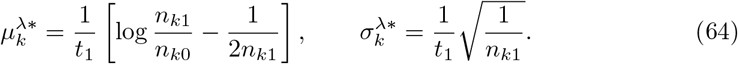

Again, compared to the maximum-likelihood estimate *λ*^ML^ in (54), the mean 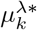 includes an entropic correction, which ensures that 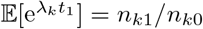 for a normally distributed *λ*_*k*_. Interestingly, the precision of the growth rate is proportional to the counts at the last time point only, while the precision of δ_*k*_ in (63) depends on the counts at both time points.

Because the standard deviations in (63) and (64) are obtained from the lower bound to the ELBO, the inferred uncertainties are expected to be smaller than those from maximizing the actual ELBO, especially for variants with high counts.

Once estimated using (63) and (64), the means can be shifted collectively to any desired gauge. In the sixth row of Table 3 (LB, *T* = 2) we show the estimates (64) calculated from the counts at only the first and third time points of the exponential dataset in (43). It is interesting to observe how the standard deviations estimated from only two time points are larger than those from all five time points, shown one line above them in Table 3 (LB, *T* = 5).

Using (64), we can calculate the means and standard deviations of the standard approach of estimating growth rates relative to a reference variant:

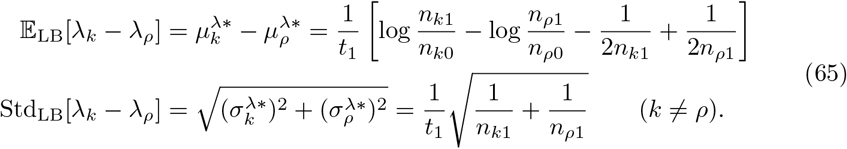

The expectation in the first line of (65) contains entropic corrections to the standard formula for growth-rate estimation (55). The new terms are expected to be numerically important in the case of very few counts. Our variational Bayesian analysis additionally provides uncertainty estimates, quantified by the standard deviations in the second line of (65). For a given variant *k*, this uncertainty is larger than that of our alternative estimator (64), because it additionally contains the counts of the reference.

This exhausts our analysis of exponential growth. In the next section, we apply the joint analysis of all sequencing time points—both in its maximum-likelihood and variational-inference versions—to non-exponential growth.

### 3.4 Beyond exponential growth: models with saturation

From a mathematical point of view, exponential growth is attractive because it admits simple linear solution. Similarly, linear least-squares regression is attractive because it can be solved by single matrix inversion. Such analytical tractability, however, is lost in the three approaches for analyzing exponential growth that we examined above—weighted least-squares fit to a *softmax* function (Sec 3.1.2), maximization of the all-time likelihood (Sec 3.2.1) or all-time ELBO (Sec 3.3) of a multinomial-*softmax* model. In these problems, a certain scalar function had to be optimized numerically with respect to either *K* or 2*K* (for ELBO) parameters. Conveniently, the gradients of the loss function could be computed through automatic differentiation [19]. In fact, the entire gradient-update procedure could be implemented with a few lines of code in PyTorch [31].

Once numerical gradient calculation becomes part of the estimation process, the analytical integrability of exponential growth loses its central importance, since the non-linear trajectories of any alternative model can be integrated numerically, while accumulating the gradients with respect to the model parameters through automatic differentiation. Leveraging this flexibility, here we apply the maximum-likelihood and variational-inference approaches to growth models with saturation. Although the strategy is illustrated for logistic and Gompertz growth, it readily extends to other growth models with a few parameters per variant.

Handling non-exponential growth is straightforward. For the log likelihood (49), we previously substituted the linear dependence of the log abundances to arrive at (50). Now we use the general growth dynamics *y*_*kτ*_ = *y*_*k*_(***λ*, δ**; *t*_*τ*_) from (35). Similarly, in the all-time ELBO (56) we now use *y*_*kτ*_ = *y*_*k*_(***µ***^*λ*^ + ***σ***^*λ*^ ⊙ ***ϵ***^*λ*^, ***µ***^δ^ + ***σ***^δ^ ⊙ ***ϵ***^δ^; *t*_*τ*_).

One important difference from exponential growth concerns the choice of the gauge. In exponential growth, this freedom exists independently for both the initial log abundances and the growth rates. In models with saturation, fixing the gauge of **δ**—by measuring abundances in units of the initial culture density, for example—also fixes the unit of the carrying capacity. Since increasing or decreasing all growth rates by an arbitrary constant would alter the time needed to reach the carrying capacity, the growth rates of the variants can no longer be shifted arbitrarily for a given 𝒦. Consequently, the values obtained by maximizing the likelihood or the ELBO correspond to absolute growth rates.

#### 3.4.1 Four variants

Maximizing the all-time log likelihood for the two models with saturation yields the estimates ***λ***^ML^ and **δ**^ML^. The variant fractions calculated from these are shown in the two right-most columns of Fig 4.

When maximizing the all-time ELBO, at each iteration step we integrated numerically *M* = 100 independent copies of the dynamics to approximate the ELBO and its gradients, as described earlier. To start the optimization of the variational parameters from reasonable values, we initialized the means with the maximum-likelihood estimates ***λ***^ML^ and **δ**^ML^, and the standard deviations according to (58). Although these latter expressions were obtained for exponential growth, they are expected to be of the right order of magnitude for the other models as well. The final variational parameters are visualized in two different ways: their Gaussian posteriors are shown in Fig 6A and the variant fractions calculated from them in Fig 6B. Results for exponential growth are also included for comparison.

**Fig 6.**
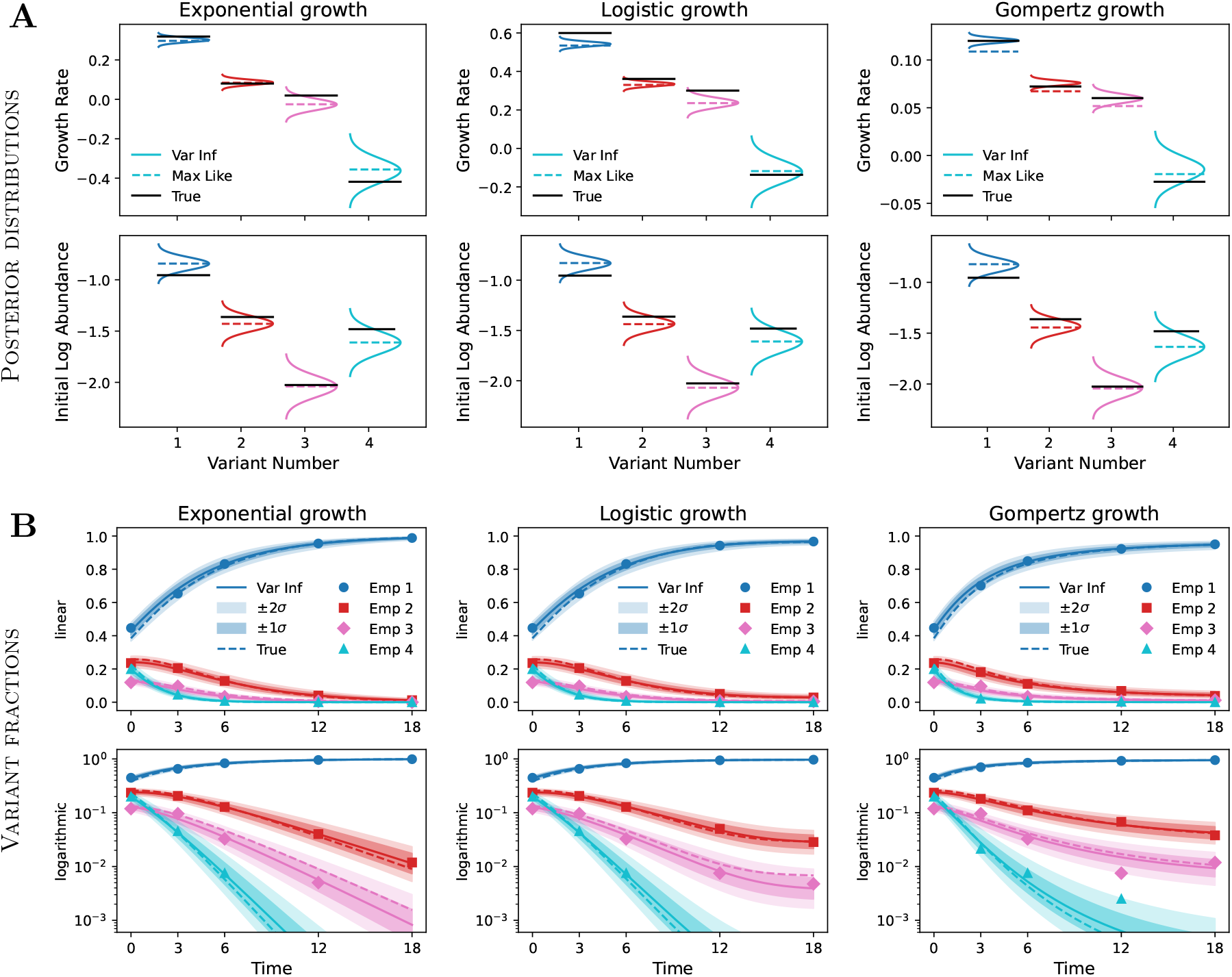
Variational inference of the parameters of the growth models. **A**: Independent Gaussian factors in the interval *µ*_*k*_ ± 3*σ*_*k*_ (bell curves, Var Inf), compared with maximum-likelihood estimates (dashed lines, Max Like) and true values (black lines, True). **B**: Time trajectories calculated from the posterior means (solid lines, Var Inf), accompanied by their uncertainty estimates (shaded bands). All true trajectories (dashed lines, True) and empirical ratios from the observed counts (symbols, Emp) lie within the *±*2*σ* confidence intervals.

When comparing the growth rates in the top row of Fig 6A across models, recall that the means are constrained to sum to zero for exponential growth, whereas no such restriction applies to logistic and Gompertz growth. In addition, both exponential and logistic growth employ the growth rates (42), while these were scaled down by a factor of five for Gompertz growth to achieve comparable saturation timescales. In contrast, the initial log abundances are treated identically across the three growth models, so their posterior Gaussian distributions in Fig 6A are directly comparable.

To facilitate direct comparison of the growth rates across models, Table 4 reverses the applied transformations: the inferred Gompertz means and standard deviations are multiplied by five, and the absolute means of the logistic and Gompertz rates are additionally shifted to a zero-sum gauge. (The corresponding values for the log abundances are given in SI, Table S2.)

**Table 4.** Variational estimates of growth rates and their uncertainties for the three growth models. Gompertz means and standard deviations are multiplied by five; true growth rates from (42).

| | $\mu^\lambda$ (absolute values) | $\mu^\lambda$ (zero-sum gauge) | $\sigma^\lambda$ (uncertainties) |
| --- | --- | --- | --- |
| Exponential | - | [0.302 0.086 -0.026 -0.362] | [0.012 0.013 0.029 0.062] |
| Logistic | [0.543 0.334 0.237 -0.121] | [0.294 0.086 -0.011 -0.370] | [0.012 0.012 0.025 0.061] |
| Gompertz $\times 5$ | [0.602 0.380 0.298 -0.073] | [0.300 0.078 -0.004 -0.375] | [0.013 0.013 0.024 0.066] |
| True $\lambda$ | [0.600 0.361 0.301 -0.137] | [0.319 0.080 +0.019 -0.418] | - |

The strong overall agreement across the three growth models observed in Fig 6 and Table 4 demonstrates that the subtle differences in the counts data (43) are properly deconvolved by the respective growth models to yield consistent parameter estimates. On a more detailed level, however, it is unclear why the means of the Gompertz growth rates deviate more strongly from their maximum-likelihood counterparts than do those of the other two models.

To compute the uncertainty bands at *±σ* and *±*2*σ* for the trajectories in Fig 6B, we generated 1000 random realizations of the standard normal variables (***ϵ***^*λ*^, ***ϵ***^δ^) and zeroed the values that were larger than, respectively, 1 and 2. Using these modified random samples, we calculated 1000 different sets of growth parameters (***λ*, δ**), and computed trajectories for each one of them. At each time point, the maximum and minimum fractions across these trajectories were used as boundaries for the confidence intervals in Fig 6B. All empirical variant fractions (symbols) and true fractions used to generate the multinomial counts data (dashed lines) lie inside the *±*2*σ* bands.

#### 3.4.2 Many variants

From a practical perspective, it is essential to evaluate how the proposed inference scheme scales to many variants. As a compromise between increasing the variant number while still allowing for visual inspection, we generated synthetic data for *K* = 100 variants under logistic growth, drawing the growth rates and initial log abundances from Gaussian distributions. The calculated variant fractions at the same five selected time points as before were used to generate multinomial counts with approximately 100*K* total counts per time point, following the procedure of Sec 2.4.4. Prior to analysis, we verified that all variants had at least two nonzero counts across the five time points. For the realization analyzed here, the numbers of variants with 2, 3, 4, and 5 nonzero counts were, respectively, 13, 37, 11, and 39 (see SI, Fig S6).

Applying the same settings as in the four-variant analysis, we obtained the Gaussian posteriors shown in Fig 7A, where the variants are ordered by their true growth rates. Overall, the estimated posterior standard deviations are larger for variants with (i) lower initial abundances and (ii) smaller growth rates, as these jointly determine the total number of counts accumulated over time. In all cases, the true parameter values (black lines) are seen to lie well within three standard deviations of the corresponding variational means.

**Fig 7.**
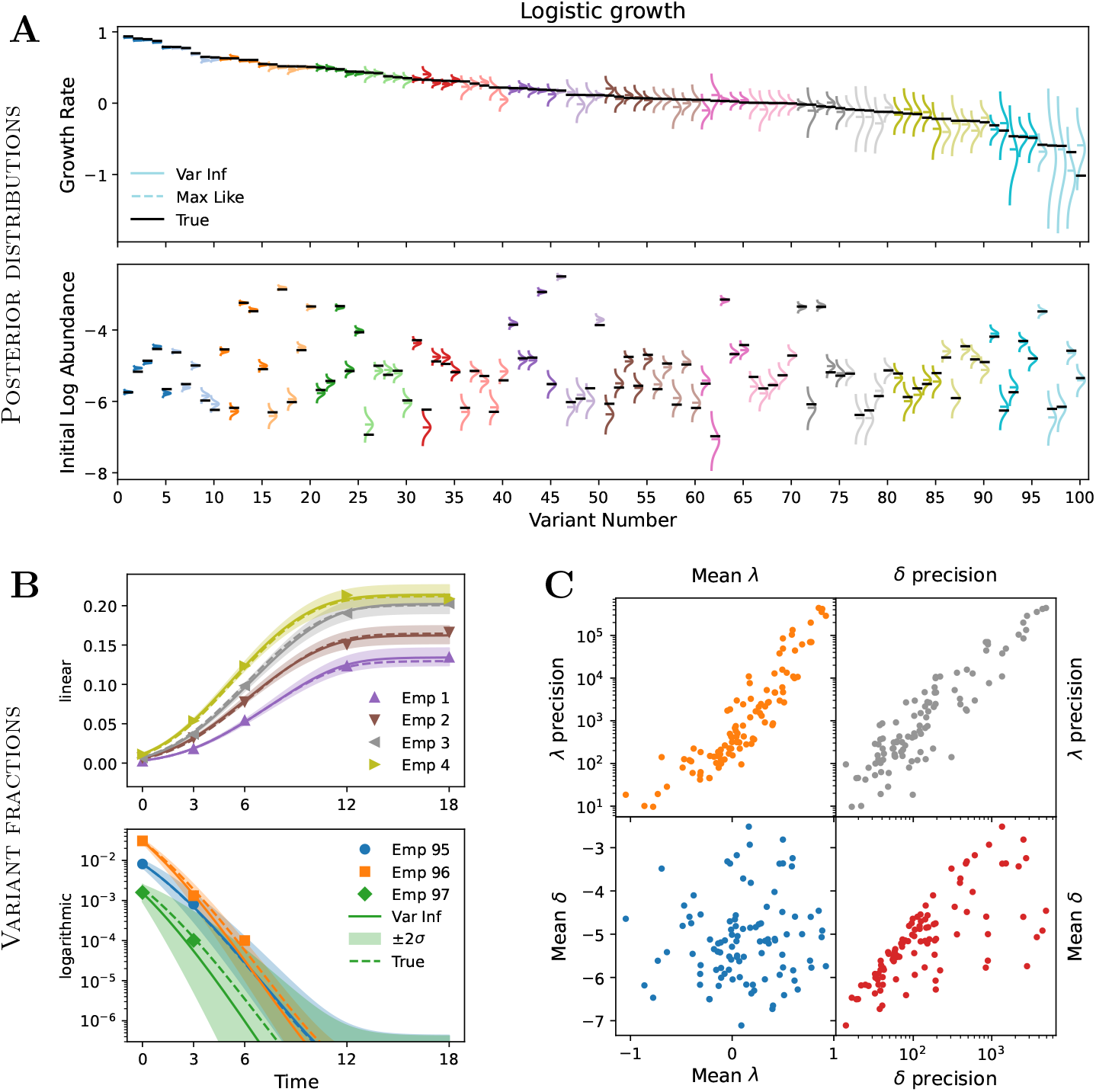
Variational inference for one hundred variants undergoing logistic growth. **A**: Gaussian posteriors (solid lines, Var Infer) with variants sorted according to their true growth rates. The true values (black lines) lie within the shown *±*3*σ* intervals from the estimated means. **B**: Time trajectories calculated using the variational parameters (solid lines, Var Inf) for the first four variants (top) and the three variants before the last three (bottom). The true trajectories (dashed lines, True) and the empirical count ratios (symbols, Emp) lie within the shown *±*2*σ* uncertainty bands. **C**: Scatter plots of the posterior means and precisions, where precision is the reciprocal variance (*λ*: growth rate; δ: initial log abundance).

Figure 7B displays the time-dependent variant fractions and their *±*2*σ* uncertainty bands for the first four variants (top panel, linear scale) and variants 95, 96 and 97 (bottom panel, logarithmic scale). In this case, the plotted confidence intervals were obtained from the maxima and minima across 5000 different trajectories. Interestingly, even the tiny growth-rate uncertainties of the fastest-growing variants 1 through 4 in Fig 7A, give rise to noticeable uncertainties of their fractions in Fig 7B. In the lower panel of Fig 7B, the uncertainty band of variant 96 (with three non-zero counts) is narrower than those of variants 95 and 97 (with two non-zero counts), in line with the number of counts of these variants.

In Fig 7C, we show the correlations between the inferred means and standard deviations of the growth parameters, with the latter expressed as precisions 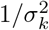. The means of the growth rates and initial log abundances in the lower-left corner confirm that these parameters were indeed drawn from independent Gaussian distributions. The upper-left corner of the plot reveals a strong positive correlation between the means and precisions of the growth rates: faster growing variants have more precisely estimated rates. A similar but weaker trend is observed for the initial log abundances (lower-right corner), where the variants with the highest δ precision have only intermediate mean δ values. The high precision of these points can be traced to the fact that they correspond to variants with high growth rates, resulting in larger total counts over time.

Finally, we note that optimizing the ELBO for 500 steps on a single CPU took 3 seconds for the logistic-growth example with *K* = 4 variants and 13 seconds for *K* = 100 variants, under identical computational settings. Further increasing the number of variants by a factor of ten to *K* = 1000, increased the runtime by approximately the same factor to 110 seconds (see SI, Fig S7). Since the logistic dynamics in this example is integrated numerically, the total number of steps taken by the Euler integrator should directly influence the optimization time. Increasing the integration time step by a factor of two, thus reducing the number of integration steps from 180 to 90, resulted in runtimes of 7.1 and 50 seconds, respectively, for *K* = 100 and *K* = 1000, which are about a factor of two smaller than the previous times of 13 and 110 seconds. For more complex variant dynamics, or when the number of variants is increased even further, using higher-order integration schemes like Heun and Runge-Kutta may be advantageous. (This consideration is not relevant for exponential growth where the dynamics are calculated analytically.)

## 4 Concluding discussion

To the best of our knowledge, the *softmax* transformation has not been applied previously to the estimation of variant growth rates from sequencing counts, despite its recognized importance in the analysis of compositional data [18, 24]. In this work, we revisited three established approaches for growth-rate estimation: least-squares regression (Sec 3.1), likelihood maximization (Sec 3.2), and variational Bayesian inference (Sec 3.3). In each case, linking the probabilistic model of sequencing counts to the deterministic model of variant growth via the *softmax* transformation yielded new analytical insights and results.

In the context of weighted least-squares regression for exponential growth [9], we argued that fitting a *softmax* -based analytical model to the data is preferable to fitting a linear model (Sec 3.1). The former better aligned instances of zero counts with the overall trends of the non-zero counts (Fig 3) and yielded estimates that were numerically invariant to the choice of reference variant (Table 2). Notably, we derived the weights of the mean-squared errors for these two alternative fitting schemes from Bayesian analysis with a Dirichlet posterior (Sec 2.2).

Despite these improvements, we contend that least-squares regression is ultimately inadequate for integrating sequencing counts across time points for two main reasons. First, the Bayesian uncertainty estimates apply to variant fractions rather than to the growth rates, which are the quantities of primary interest. Second, temporal integration is handled more naturally and rigorously by maximizing the likelihood under an appropriate noise model.

Following Razo-Mejia and colleagues [11], here we adopted a multinomial model of noise, because it inherently respects the compositional constraint that variant fractions must sum to unity [18], capturing the negative covariances among them—an increase in the fraction of one variant necessarily decreases the fractions of the others. In contrast, modeling the stochasticity of individual variant counts using Poisson [8] or negative-binomial [10] distributions neglects this essential compositional structure.

Using the *softmax* function, we transformed the standard parameters of the multinomial distribution—variant fractions—into log abundances, which parametrized the resulting multinomial-*softmax* likelihood (Sec 3.2.1). While our motivation was to incorporate the growth-model parameters directly into the likelihood, the *softmax* transformation, in fact, naturally arises when the multinomial distribution is expressed as a member of the *exponential family* of distributions, where the new parameters—our log abundances—become the *natural parameters* of the distribution [21].

In general, the maximization of the multinomial-*softmax* likelihood across multiple time points cannot be performed analytically even for exponential growth, where the log abundances are linear in time (Sec 3.2.1). An exception arises in the special case of sequencing at only two time points (Sec 3.2.2). In this experimentally attractive scenario, the *softmax* perspective led to a growth-rate estimator that uses the counts of a single variant [Eq (54)]. In contrast, the standard growth-rate estimator [Eq (55)] additionally compares with the counts of a reference variant. In this latter case, the growth rate of the reference relative to itself is, by definition, zero, which makes the associated uncertainty ill-defined. Since our alternative growth-rate estimator does not compare with a reference variant, it does not suffer from this ambiguity.

The final methodology we examined through the lens of the *softmax* transformation (Sec 3.3) was the Bayesian framework of Razo-Mejia and colleagues [11], in which variational inference is used to estimate growth rates and quantify their associated uncertainties. In the case of exponential growth, expressing the variant fractions as a *softmax* transformation of the linear log abundances eliminated the need to first estimate the mean growth rate of the population (see SI, Sec S1), thereby simplifying the procedure of Ref [11].

One disconcerting aspect of variational inference with Gaussian posterior is the rapid proliferation of parameters that must be inferred from the data. When applied to a single sequencing time point, where growth dynamics are irrelevant, the Gaussian posterior introduced twice as many parameters as the Dirichlet posterior—and, in fact, twice as many as the number of observed counts (Sec 2.4). We therefore examined how maximizing the ELBO determined 2*K* parameters from only *K* experimental observations and clarified the distinct roles of the posterior means and standard deviations: the means capture fractional information appropriate for compositional data, whereas the standard deviations encode information about absolute count magnitudes, which ultimately dictate the uncertainty of the estimated parameters (Sec 2.4.1). This observation was further supported by the analytical lower bound to the ELBO, derived via Jensen’s inequality, whose maximization provided closed-form expressions for the optimal parameters (Sec 2.4.2).

Just like the likelihood function on which it is based, the ELBO [Eq (56)] naturally integrates data across multiple sequencing time points. However, because the expectation over the posterior distribution, which is contained within the ELBO, cannot be computed analytically, only limited insight into the ELBO optimum can be obtained from examining its gradients. Even the analytical lower bound to the ELBO, derived from Jensen’s inequality [Eq (59)], could not be maximized in closed form when data from several time points were integrated—except in the special case of only two time points.

In this simpler case, we maximized the ELBO lower bound analytically, and expressed the means and standard deviations in terms of the variant counts at the two time points (Sec 3.3.2). To our knowledge, the resulting growth-rate estimator [Eq (64)] has not been reported previously. Its inferred variance scales inversely with the counts at the second time point only, and introduces an entropic correction to the mean compared to the maximum-likelihood estimator (54). For the illustrative numerical example with four variants, the uncertainties estimated by maximizing the lower bound underestimated those assigned by maximizing the ELBO, particularly for the variant with large counts (Fig 5).

The *softmax* transformation provided a natural interface between variant fractions, which parameterize the multinomial model of counting noise, and variant log abundances, which evolved linearly with time under exponential growth. At the same time, this interface delineated a clear conceptual boundary between the probabilistic and deterministic components that are jointly needed to analyze sequencing data across multiple time points. This separation highlighted that the log abundances—the inputs to the *softmax* function—could, in principle, originate from any underlying model of growth (Sec 2.5). Building on this observation, we extended the maximum-likelihood and variational-inference frameworks to logistic and Gompertz dynamics (Sec 3.4). In this context, the use of automatic differentiation [19, 32] was indispensable, as it allowed gradients of the objective function to be accumulated seamlessly during the numerical integration of the growth model.

The three growth models examined here were parameterized directly in terms of variant-specific growth rates. However, more mechanistic formulations—in which the growth rates emerge as a function of underlying microscopic parameters—are equally conceivable. For instance, in studies of resistance to beta-lactam antibiotics, models have been proposed that express the growth rate of a variant in terms of the biochemical parameters *V*_max_ and *K*_*M*_ of the beta-lactamase enzyme it encodes [16, 17]. Incorporating such models into the present inference framework could enable the simultaneous estimation of these parameters and their associated uncertainties.

The ability to incorporate arbitrary growth models into the proposed inference framework will open exciting opportunities for high-throughput inference of diverse biochemical parameters that influence growth, through sequencing-based measurements of thousands of variants in pooled competition assays. The work presented here is a first step toward this goal.

## Supporting information

**S1 Text. Supporting information**. Contains some of the mathematical detail that is not shown in the main text and several additional figures.

## Acknowledgments

We thank all Toprak Lab members—Fatma Coskun, Ilona Gaszek, Adam Lyon, and Sadık Yıldız—for fostering an inspiring research environment. DS is grateful to Kim Reynolds, Phil Brown, and Jerry Dinan for offering an excellent introductory course at UTSW on pooled competition experiments, through which he became aware of the Enrich2 software tool [9]. Their interest in applying the Bayesian framework of Ref [11] to data analysis provided strong initial motivation for this work. DS also extends heartfelt gratitude to Ali Rana Atılgan for his constant encouragement and insightful feedback on this manuscript.

