## Supporting information for "Counting-based inference of mutant growth rates from pooled sequencing across growth regimes"

Deniz Sezer*[1] and Erdal Toprak[1]

[1]Department of Pharmacology, UTSW Medical Center, TX 75390, USA

November 16, 2025

#### S1 Describing exponential growth

We denote the total number of different variants in the pooled competition assay by $K$, and index the individual variant types by $k = 1, \ldots, K$. When modeling the time dependence of the variants, we assume that each cell belongs to a single variant, so variant abundances increase or decrease in proportion to the total number of their corresponding cells. We model the change in cell numbers through a deterministic model of growth.

In exponential growth, the number of cells of variant $k$ at time $t$, denoted by $N_k(t)$, change in time according to the following differential equation:

$$\frac{\mathrm{d}N_k(t)}{\mathrm{d}t} = \lambda_k N_k(t). \tag{S1}$$

Here, $\lambda_k$ is the specific growth rate of variant $k$, which is modeled as being constant throughout the experiment. Note that the dynamics of one variant is independent from the dynamics of the other variants.

When modeling the stochasticity of the sequencing read counts with some probability distribution, the variant fractions are the natural parameters in terms of which the distribution is described analytically. With

$$N(t) = \sum_{k=1}^{K} N_k(t) \tag{S2}$$

denoting the total number of cells across all $K$ variants, the fraction of variant $k$ is

$$f_k(t) = \frac{N_k(t)}{N(t)}. \tag{S3}$$

We now derive a differential equation describing how $f_k(t)$ changes in time.

Starting with the total number of cells in the denominator of (S3), we differentiate (S2) with respect to time and use (S1) to obtain

$$\frac{\mathrm{d}N(t)}{\mathrm{d}t} = \sum_{k=1}^{K} \lambda_k N_k(t) = \bar{\lambda}(t) N(t). \tag{S4}$$

*

In the second equality, we introduced the instantaneous mean growth rate of the population

$$\bar{\lambda}(t) = \sum_{k=1}^{K} \lambda_k f_k(t). \tag{S5}$$

When the variant fractions change with time, this mean growth rate also changes, even if the individual growth rates of the variants remain constant.

Using (S1) and (S4), we can now differentiate the definition (S3) with respect to time to get

$$\frac{\mathrm{d}f_k(t)}{\mathrm{d}t} = \frac{1}{N(t)}\frac{\mathrm{d}N_k(t)}{\mathrm{d}t} - \frac{N_k(t)}{N(t)}\frac{1}{N(t)}\frac{\mathrm{d}N(t)}{\mathrm{d}t} = f_k(t)\lambda_k - f_k(t)\bar{\lambda}(t). \tag{S6}$$

Hence,

$$\frac{\mathrm{d}f_k(t)}{\mathrm{d}t} = [\lambda_k - \bar{\lambda}(t)]f_k(t). \tag{S7}$$

Importantly, the instantaneous rate of change of variant $k$ (in the square brackets) contains the mean growth rate of the population, which depends on the fractions of all other variants. Hence, when described in terms of the variant fractions, the variant dynamics is neither exponential nor decoupled.

Because the variant fractions parametrize the probability distribution of the sequencing counts, there is a tendency in the literature to also describe the growth dynamics in terms of them.[1;2;3;4] Specifically, the differential equation (S7) is commonly integrated between two successive sequencing time points as[3]

$$f_k(t_{\tau+1}) = f_k(t_\tau)\mathrm{e}^{[\lambda_k - \bar{\lambda}(t_\tau)](t_{\tau+1}-t_\tau)}, \tag{S8}$$

where $t_\tau$, with $\tau = 0, 1, \ldots, T-1$, are the time points of sequencing. Note that (S8) is an approximation to the true dynamics, since the integration from $t_\tau$ to $t_{\tau+1}$ assumes that the mean growth rate remains constant in the time interval between two successive sequencing time points.

When estimating variant growth rates using (S8), it becomes necessary to first estimate the mean growth rate $\bar{\lambda}(t_\tau)$, before estimating the individual growth rates.[1;2;3;4] In evolution experiments, where the spontaneously occurring mutants initially comprise a tiny fraction of the population, the mean growth rate can be safely identified with the growth rate of the wild type.[1] However, such an approximation is not justified in growth competition experiments where all variants may be present from the start at similar abundances. One workaround, which effectively recreates the situation in evolution experiments, is to ensure that the wild type constitutes at least 95% of the population throughout the duration of the competition experiment.[3;4] Biologically, this also ensures that all other low-abundance variants grow in a wild-type context. It is important, however, to distinguish this biological requirement from the mathematical condition that the mean growth rate is approximately constant. While the wild-type context could be desired in itself, we argue that the mathematical approximation is not needed for the purposes of estimating growth rates.

Our approach is to describe exponential growth not directly in terms of variant fractions but in terms of the natural logarithms of the cell counts, which we call *log abundances*:

$$y_k(t) = \log N_k(t) \tag{S9}$$

These *log abundances* are better suited than the variant fractions for describing exponential growth, as we show next.

Differentiating the definition (S9) with respect to time, and using (S1), we obtain

$$\frac{\mathrm{d}y_k(t)}{\mathrm{d}t} = \frac{1}{N_k(t)}\frac{\mathrm{d}N_k(t)}{\mathrm{d}t} = \lambda_k. \tag{S10}$$

This differential equation can now be integrated exactly (i.e., without assuming that the mean growth rate of the population is constant in the integration interval). The result is

$$y_k(t) = \lambda_k t + \delta_k, \tag{S11}$$

where the integration constant

$$\delta_k = \log N_k(0) \tag{S12}$$

reflects the abundance of variant $k$ at the beginning of the competition experiment.

For compact notation, we write variant-specific properties as $K$-dimensional (column) vectors in bold. For example, $\boldsymbol{\lambda} = (\lambda_1, \ldots, \lambda_K)$ collects the growth rates of all variants. For time-dependent quantities, we denote evaluation at the sequencing times $t_\tau$ with a subscript, e.g., $f_{k\tau} = f_k(t_\tau)$, and in vector form $\mathbf{f}_\tau = (f_{1\tau}, \ldots, f_{K\tau})$.

At this stage, we have two complementary representations: *variant fractions*, which naturally parametrize the probabilistic noise model and can therefore be directly estimated from sequencing counts, and *variant log abundances*, which conveniently describe the growth dynamics through their linear dependence on the growth parameters. These two descriptors are straightforwardly linked by combining their definitions in (S3) and (S9):

$$f_k(t) = \frac{\mathrm{e}^{y_k(t)}}{\sum_{i=1}^{K} \mathrm{e}^{y_i(t)}} = \phi_k(\mathbf{y}(t)), \tag{S13}$$

where the mapping $\boldsymbol{\phi}(\cdot)$ is known as the *softmax* transformation.[5] Note that this relationship between fractions and log abundances is an identity which follows from the definitions of $f_k(t)$ and $y_k(t)$, and thus holds independently of the growth dynamics.

In the case of exponential growth,

$$f_k(t) = \frac{\mathrm{e}^{\lambda_k t + \delta_k}}{\sum_{i=1}^{K} \mathrm{e}^{\lambda_i t + \delta_i}}. \tag{S14}$$

Unlike (S8), the time dependence of the variant fractions in (S14) is exact. Moreover, because the mean growth rate does not appear explicitly, the individual growth rates can be estimated directly, without the need to first estimate the mean growth rate of the culture.

In summary, the *softmax* transformation (S13) provides a bridge between the intuitive parameters of the probabilistic noise model (fractions on the left) and the mathematically convenient description of biological growth (log abundances on the right).

### S2 Independent treatment of the counting noise at each sequencing time point

In this section, we provide derivations for the equations in Secs. 2.1 and 2.2 of the main text. Rather than showing these standard derivations in their most general form, we present them in the specific context of interest to us, namely, the analysis of the read counts at one sequencing time point. In the first subsection, we maximize the likelihood of the multinomial distribution and in the second subsection, we move on to Bayesian inference with a conjugate Dirichlet posterior. While we explicitly show the time index when maximizing the likelihood, we stop doing so when discussing Bayesian inference in order to declutter the notation.

#### S2.1 Maximum likelihood

Here we estimate the parameters of the multinomial distribution by maximizing its likelihood for given counts data. We do this first for the standard parametrization of the multinomial distribution and then again for its reparametrization through the *softmax* transformation.

##### S2.1.1 Working with fractions

The multinomial distribution assigns probabilities to the possible combinations of variant read counts given the proportions of these variants. Denoting the variant counts by the $K$-component counts vector $\mathbf{n}$, and their fractions by the vector $\mathbf{f}$, the assigned probabilities are

$$p(\mathbf{n}|\mathbf{f}) = \frac{1}{C(\mathbf{n})} \prod_{k=1}^{K} (f_k)^{n_k}. \tag{S15}$$

The normalization factor

$$C(\mathbf{n}) = \frac{1}{n!} \prod_{k=1}^{K} (n_k!) \tag{S16}$$

ensures that the sum of these probabilities over all possible count combinations equals one. Being the actual fractions of the variants, the parameters of this model satisfy the conditions

$$0 \leq f_k \leq 1, \qquad \sum_{k=1}^{K} f_k = 1. \tag{S17}$$

The probability in (S15), which assigns statistical weights to the counts data, can be viewed as the probabilistic rule according to which the counts data is generated. From that perspective, it constitutes our generative model. Our purpose is to estimate the parameters of the model from counts data that has been generated from it. We do this by maximizing the probability in (S15) with respect to the parameters $\mathbf{f}$ for given counts. When viewed as a function of the parameters, the probability is referred to as a *likelihood* and hence this estimation strategy is called *maximum likelihood*.

At a given sequencing time $t_\tau$, the logarithm of the likelihood is

$$\mathcal{L}(\mathbf{f}_\tau|\mathbf{n}_\tau) = \log p(\mathbf{n}_\tau|\mathbf{f}_\tau) = \sum_{k=1}^{K} n_{k\tau} \log f_{k\tau} + \text{const}, \tag{S18}$$

where $\mathbf{n}_\tau$ and $\mathbf{f}_\tau$ are, respectively, the sequencing read counts and the true fractions of the variants at time $t_\tau$. We have only shown the part of the log likelihood that depends on the parameters, and have put all other terms into the constant, which becomes irrelevant when maximizing with respect to the parameters.

Because their sum equals one, only $K-1$ of the $K$ fractionparameters are linearly independent. To maximize the log likelihood while preserving the sum condition in (S17), we maximize the auxiliary function

$$\mathcal{H}(\mathbf{f}_\tau, \nu|\mathbf{n}_\tau) = \mathcal{L}(\mathbf{f}_\tau|\mathbf{n}_\tau) - \nu_\tau \left( \sum_{k=1}^{K} f_{k\tau} - 1 \right), \tag{S19}$$

where $\nu_\tau$ is a Lagrange multiplier. The partial derivative of this function with respect to $f_{k\tau}$ is

$$\frac{\partial \mathcal{H}}{\partial f_{k\tau}} = \frac{n_{k\tau}}{f_{k\tau}} - \nu_\tau. \tag{S20}$$

The maximum-likelihood estimates of the variant fractions are obtained by equating this derivative to zero, hence

$$n_{k\tau} = \nu_\tau f_{k\tau}^{\text{ML}}. \tag{S21}$$

Summing this last result over all variants and using (S17), we find that the Lagrange multiplier equals the total number of sequencing reads at this time point:

$$\nu_\tau = \sum_{k=1}^{K} n_{k\tau} = n_\tau. \tag{S22}$$

We can now eliminate the Lagrange multiplier from (S21) to arrive at

$$f_{k\tau}^{\text{ML}} = \frac{n_{k\tau}}{n_\tau}, \tag{S23}$$

which is the result reported in Eq (6) of the main text. Thus, by calculating the fraction of the reads of variant $k$ among all reads, we in fact calculate the maximum-likelihood estimates of the true variant fractions that parametrize the multinomial distribution.

##### S2.1.2 Working with log abundances

Here we switch from the variant fractions to the log abundances using (S13). Since the fractions $f_{k\tau}$ obtained from the *softmax* transformation automatically satisfy the conditions in (S17), there is no need for a Lagrange multiplier during the maximization of the log likelihood

$$\mathcal{L}(\mathbf{y}_\tau|\mathbf{n}_\tau) = \sum_{k=1}^{K} n_{k\tau} y_{k\tau} - n_\tau \log\left(\sum_{k=1}^{K} \mathrm{e}^{y_{k\tau}}\right). \tag{S24}$$

The derivatives with respect to the new parameters are

$$\frac{\partial \mathcal{L}}{\partial y_{k\tau}} = n_{k\tau} - n_\tau \frac{\mathrm{e}^{y_{k\tau}}}{\sum_{i=1}^{K} \mathrm{e}^{y_{i\tau}}} \tag{S25}$$

and so the maximum-likelihood estimates of the log abundances satisfy the equality

$$\phi_k(\mathbf{y}_\tau^{\mathrm{ML}}) = \frac{\mathrm{e}^{y_{k\tau}^{\mathrm{ML}}}}{\sum_{i=1}^{K} \mathrm{e}^{y_{i\tau}^{\mathrm{ML}}}} = \frac{n_{k\tau}}{n_\tau}. \tag{S26}$$

This result is given as eq. (15) in the main text.

While (S26) is consistent with (S23), the latter is a closed-form expression for the maximum-likelihood fractions. In contrast, the equalities (S26) still have to be solved for the ML log abundances, which requires the inversion of the *softmax* transformation. However, the *softmax* function does not have a unique inverse because it is invariant to a common shift of all components of its argument.

Taking the logarithm of both sides of (S26), we obtain

$$y_{k\tau}^{\mathrm{ML}} - \sum_{i=1}^{K} \mathrm{e}^{y_{i\tau}^{\mathrm{ML}}} = \log n_{k\tau} - \log n_\tau. \tag{S27}$$

Up to an arbitrary, variant-independent shift, therefore, the maximum-likelihood estimates of the log abundances are given by

$$y_{k\tau}^{\mathrm{ML}} = \log n_{k\tau} + c_\tau, \tag{S28}$$

where the constant $c_\tau$ is the same across the variants but can change at different time points.

#### S2.2 Bayesian analysis for fractions

In this subsection, we continue to analyze the different sequencing time points separately from each other, but suppress the time superscript.

All Bayesian analysis, either exact or approximate, makes use of the Bayes equality

$$p(\mathbf{f}|\mathbf{n}) = \frac{p(\mathbf{n}|\mathbf{f})p(\mathbf{f})}{p(\mathbf{n})}, \tag{S29}$$

which we have written here in the context of counts, $\mathbf{n}$, that are generated from a multinomial distribution with parameters $\mathbf{f}$. The purpose in Bayesian inference is to obtain the posterior distribution of the parameters given the data, which is on the left-hand side of (S29), starting from a prior distribution and a likelihood, which are in the numerator on the right-hand side of (S29). The denominator in (S29) is

$$p(\mathbf{n}) = \int p(\mathbf{n}|\mathbf{f})p(\mathbf{f})\mathrm{d}\mathbf{f} \tag{S30}$$

and ensures that the posterior is properly normalized with respect to $\mathbf{f}$.

When the likelihood in (S29) is a multinomial distribution, it is convenient to select the prior to be a Dirichlet distribution, since the the posterior is also Dirichlet but with some other parameters.

The probability density function of a Dirichlet distribution with parameters $\boldsymbol{\alpha}$ is

$$p(\mathbf{f}|\boldsymbol{\alpha}) = \frac{1}{B(\boldsymbol{\alpha})}\prod_{k=1}^{K}(f_k)^{\alpha_k-1}. \tag{S31}$$

The normalization factor,

$$B(\boldsymbol{\alpha}) = \frac{1}{\Gamma(\alpha)}\prod_{k=1}^{K}\Gamma(\alpha_k), \tag{S32}$$

involves the gamma function, $\Gamma(\cdot)$, and we have denoted the sum of the parameters over all variants by

$$\alpha = \sum_{k=1}^{K}\alpha_k. \tag{S33}$$

The parameters of the Dirichlet distribution should be positive but, unlike the fractions $\mathbf{f}$, they need not sum to one. Thus, the Dirichlet distribution uses $K$ linearly independent numbers (the parameters $\boldsymbol{\alpha}$) to describe the distribution of $K-1$ linearly independent numbers (the fractions $\mathbf{f}$).

To show that the posterior is also Dirichlet, we first take the logarithms of both sides of the Bayes rule (S29):

$$\log p(\mathbf{f}|\mathbf{n}) = \log p(\mathbf{n}|\mathbf{f}) + \log p(\mathbf{f}) + \text{conts}. \tag{S34}$$

Since we focus on the dependence of the posterior on $\mathbf{f}$, all terms that do not depend on these variables have been lumped into the constant on the right-hand side of (S34). Substituting the multinomial distribution from (S15) into (S34), we obtain

$$\log p(\mathbf{f}|\mathbf{n}) = \sum_{k=1}^{K} n_k \log f_k + \log p(\mathbf{f}) + \text{const}. \tag{S35}$$

Now we take the prior to be a Dirichlet distribution with parameters $\tilde{\boldsymbol{\alpha}}$. Because the Dirichlet distribution is a conjugate prior of the multinomial distribution, the posterior is also going to be Dirichlet with some parameters, which we denote by $\boldsymbol{\alpha}$. Substituting the log of (S31) into (S35), both as a posterior and as a prior, we arrive at

$$\sum_{k=1}^{K}(\alpha_k - 1)\log f_k = \sum_{k=1}^{K} n_k \log f_k + \sum_{k=1}^{K}(\tilde{\alpha}_k - 1)\log f_k + \text{const}. \tag{S36}$$

From here, matching the coefficient of $\log f_k$, we conclude that

$$\alpha_k = n_k + \tilde{\alpha}_k, \tag{S37}$$

which is eq. (8) in the main text.

Since, in general, the variant counts are going to be different at different sequencing time points, the posterior parameters are also going to be different. However, the same prior parameters $\tilde{\boldsymbol{\alpha}}$ can be used for all time points. Furthermore, one can use the same prior parameter for all variants, i.e., $\tilde{\alpha}_k = \tilde{\alpha}$ independently of $k$. Two choices stand out:

$$\tilde{\alpha} = \begin{cases} 1, & \text{uniform prior} \\ \frac{1}{2}, & \text{Jeffreys prior.} \end{cases} \tag{S38}$$

The second option is used in Enrich2, for example.[6] We use the first option in our numerical examples.

Once the parameters of the posterior distribution are obtained, they can be used to calculate some descriptive statistics, like the means and variances of the variant fractions. For a Dirichlet distribution with parameters $\boldsymbol{\alpha}$ these are known to be

$$\bar{f}_k = \mathbb{E}_{\text{Dir}}[f_k] = \frac{\alpha_k}{\alpha}, \qquad \text{Var}_{\text{Dir}}[f_k] = \frac{\bar{f}_k(1-\bar{f}_k)}{\alpha+1}. \tag{S39}$$

In the linear least-squares fit, we also use the means and (co)variances for the logarithms of the fractions:

$$\mathbb{E}_{\text{Dir}}[\log f_k] = \psi_0(\alpha_k) - \psi_0(\alpha), \quad \text{Var}_{\text{Dir}}[\log f_k] = \psi_1(\alpha_k) - \psi_1(\alpha), \quad \text{Cov}_{\text{Dir}}[\log f_k, \log f_j] = -\psi_1(\alpha), \tag{S40}$$

where $\psi_0(\cdot)$ and $\psi_1(\cdot)$ are the digamma and the trigamma functions, respectively, and $k \neq j$ for the covariances. From these, it follows that

$$\mathbb{E}_{\text{Dir}}[\log f_i - \log f_j] = \psi_0(\alpha_i) - \psi_0(\alpha_j), \qquad \text{Var}_{\text{Dir}}[\log f_i - \log f_j] = \psi_1(\alpha_i) + \psi_1(\alpha_j), \tag{S41}$$

which appear as eq. (34) in the main text.

The digamma and trigamma functions have the following asymptotic expansions

$$\psi_0(z) \sim \log z - \frac{1}{2z} + \ldots, \qquad \psi_1(z) \sim \frac{1}{z} + \frac{1}{2z^2} + \ldots, \tag{S42}$$

whose partial sums become increasingly more accurate as the argument $z$ increases. Retaining the first two terms in the expansion of $\psi_0(\cdot)$, for example, we obtain the approximation

$$\mathbb{E}_{\text{Dir}}[\log f_i - \log f_j] \approx \left(\log \alpha_i - \frac{1}{2\alpha_i}\right) - \left(\log \alpha_j - \frac{1}{2\alpha_j}\right). \tag{S43}$$

Similarly, retaining only the first term in the asymptotic expansion of $\psi_1(\cdot)$, we get

$$\text{Var}_{\text{Dir}}[\log f_i - \log f_j] \approx \frac{1}{\alpha_i} + \frac{1}{\alpha_j}. \tag{S44}$$

### S3 Variational inference for log abundances

To introduce variational inference, we set out to carry the same Bayesian analysis as in Sec. S2.2, but for the multinomial distribution that has been parametrized in terms of the log abundances, rather than the fractions. In other words, we pursue the inference of the Bayesian posterior $p(\boldsymbol{y}_\tau|\mathbf{n}_\tau)$, which is the probability of the log abundances at time $t_\tau$ given the counts data at this time. Because this posterior cannot be obtained in closed form, unlike the Dirichlet posterior of Sec. S2.2, we determine it approximately using the framework of *variational inference* outlined in the current section.

Suppressing the time index, as we did earlier in Sec. S2.2, we write the posterior of the new parameters as

$$p(\mathbf{y}|\mathbf{n}) = \frac{p(\mathbf{n}|\mathbf{y})p(\mathbf{y})}{p(\mathbf{n})}, \tag{S45}$$

where the generative model $p(\mathbf{n}|\mathbf{y})$ refers to the multinomial distribution but with parameters obtained from a *softmax* transformation of the log abundances $\mathbf{y}$. As before, we want to obtain this posterior by combining the generative model with some reasonable prior. Since in this case it is not possible to write down the posterior distribution analytically, the strategy is to approximate it by some analytical distribution function $q_\theta(\mathbf{y})$. The idea is to select the values of the parameters $\theta$ such that the approximating function is closest to the true (but unknown) posterior.

A measure of closeness between two probability distributions that is commonly used in this type of analysis is the Kullback-Leibler (KL) divergence. Specifically, the KL divergence between the exact posterior and its approximation is

$$D_{\text{KL}}(q_\theta(\mathbf{y})||p(\mathbf{y}|\mathbf{n})) = \mathbb{E}_{\mathbf{y}\sim q_\theta}\left[\log \frac{q_\theta(\mathbf{y})}{p(\mathbf{y}|\mathbf{n})}\right], \tag{S46}$$

where $\mathbb{E}_{\mathbf{y}\sim q_\theta}[\cdot]$ denotes expectation with respect to the random variables $\mathbf{y}$ that are drawn from the distribution $q_\theta$. Importantly, this divergence is non-negative, i.e.,

$$D_{\text{KL}}(q_\theta(\mathbf{y})||p(\mathbf{y}|\mathbf{n})) \geq 0, \tag{S47}$$

with the equality being satisfied only when the approximating distribution is identical to the true posterior.

Because of the expectation over the latent random variables $\mathbf{y}$, the KL divergence in (S46) is a function only of the variational parameters $\theta$ and the data $\mathbf{n}$:

$$\mathcal{D}(\theta|\mathbf{n}) := D_{KL}(q_\theta(\mathbf{y})||p(\mathbf{y}|\mathbf{n})). \tag{S48}$$

From the perspective of the KL divergence, the best approximation $q_\theta$ to the exact posterior that we can obtain is the one whose parameters $\theta$ minimize the KL divergence, ideally, reducing it all the way down to zero. Hence, we can select the parameters of the approximating functions as

$$\theta^* = \arg\min_\theta \mathcal{D}(\theta|\mathbf{n}). \tag{S49}$$

This optimization problem is the central tool for obtaining a variational approximation of the posterior.

#### S3.1 Evidence lower bound

The main issue with the minimization in (S49) is that the KL divergence depends on the posterior distribution, which is unknown. This is in fact the distribution that we are trying to approximate because it cannot be calculated exactly. To turn (S49) into an optimization problem that can be carried out in practice, we use the Bayes rule to replace the true but unknown posterior by the right-hand side of (S45). Substituting into the definition of the KL divergence in (S46), we obtain

$$\mathcal{D}(\theta|\mathbf{n}) = \mathbb{E}_{\mathbf{y}\sim q_\theta}\left[\log \frac{q_\theta(\mathbf{y})}{\frac{p(\mathbf{n}|\mathbf{y})p(\mathbf{y})}{p(\mathbf{n})}}\right] = \mathbb{E}_{\mathbf{y}\sim q_\theta}\left[\log \frac{q_\theta(\mathbf{y})}{p(\mathbf{n}|\mathbf{y})p(\mathbf{y})}\right] + \log p(\mathbf{n}). \tag{S50}$$

When writing the second equality, we used the fact that $\log p(\mathbf{n})$ does not depend on the latent random variables $\mathbf{y}$ and is thus not affected by the expectation. Now we define

$$\mathcal{F}(\theta|\mathbf{n}) = \mathbb{E}_{\mathbf{y}\sim q_\theta}\left[\log \frac{p(\mathbf{n}|\mathbf{y})p(\mathbf{y})}{q_\theta(\mathbf{y})}\right] = \mathbb{E}_{\mathbf{y}\sim q_\theta}\left[\log p(\mathbf{n}|\mathbf{y})\right] - \mathbb{E}_{\mathbf{y}\sim q_\theta}\left[\log q_\theta(\mathbf{y})\right] + \mathbb{E}_{\mathbf{y}\sim q_\theta}\left[\log p(\mathbf{y})\right] \tag{S51}$$

and rewrite (S50) as

$$\mathcal{D}(\theta|\mathbf{n}) + \mathcal{F}(\theta|\mathbf{n}) = \log p(\mathbf{n}). \tag{S52}$$

This expression forms the basis for replacing the optimization in (S49), which is impossible to perform in practice, by an optimization that can actually be carried out numerically, as we explain now.

Because the right-hand side of (S52) does not depend on the parameters $\theta$ and, additionally, the KL divergence on the left-hand side is non-negative, *minimizing* the KL divergence with respect to $\theta$ is equivalent to *maximizing* the function $\mathcal{F}(\theta|\mathbf{n})$. When the KL divergence approaches zero, $\mathcal{F}(\theta|\mathbf{n})$ approaches $\log p(\mathbf{n})$ from below, as seen from (S52). Since $\mathcal{F}(\theta|\mathbf{n})$ is a lower bound to $\log p(\mathbf{n})$, it is commonly called *evidence lower bound* (ELBO), with *evidence* referring to the logarithm of the marginalized likelihood $p(\mathbf{n})$ (marginalized over the latent variables). Thus, the minimization of the KL divergence in (S49) is equivalent to the following maximization of the ELBO:

$$\theta^* = \arg\max_\theta \mathcal{F}(\theta|\mathbf{n}). \tag{S53}$$

Differently from the KL divergence, the ELBO can be actually calculated because all three probability distributions on the right-hand side of (S51)—the generative model, the approximating function and the prior—are known functions.

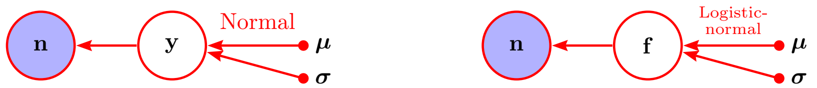


Figure S1: Approximation of the distribution of the latent random variables $y_k$ by independent Gaussians with means $\mu_k$ and standard deviations $\sigma_k$ (left). When expressed in terms of the latent variables $\mathbf{f}$ , this model involves the logistic-normal distribution of Aitchison.[7] but with a diagonal covariance matrix (right)

#### S3.2 Approximation by Gaussians

A common and convenient choice of approximating distribution $q_\theta$, with respect to which the expectations in (S51) need to be calculated, is the normal distribution with Gaussian distribution function. For a $K$-dimensional latent variable $\mathbf{y}$, the most general multivariate Gaussian is

$$q_\theta(\mathbf{y}) = \sqrt{\frac{\det(\mathbf{\Sigma})}{(2\pi)^K}}\, \mathrm{e}^{-\frac{1}{2}\sum_{ij}(y_i-\mu_i)\Sigma_{ij}(y_j-\mu_j)}, \qquad \theta = \{\boldsymbol{\mu}, \mathbf{\Sigma}\}, \tag{S54}$$

where $\boldsymbol{\mu}$ is a vector of means and $\mathbf{\Sigma}^{-1}$ is a $K \times K$ covariance matrix. To estimate this distribution from the data one would need to estimate $K + K(K+1)/2$ parameters in total.

##### S3.2.1 Mean-field approximation

Because the number of parameters increases quadratically with the number of variants and we are interested in modeling data from thousands of variants, we drastically simplify the approximating function by assuming that the latent variables are not correlated with each other. Then, instead of the multivariate Gaussian distribution in (S54), we look for an approximating distribution that is a product of independent univariate Gaussians:

$$q_\theta(\mathbf{y}) = \prod_{k=1}^{K} \frac{1}{\sqrt{2\pi\sigma_k^2}}\, \mathrm{e}^{-\frac{1}{2\sigma_k^2}(y_k-\mu_k)^2}. \tag{S55}$$

In other words, the log abundance of each variant is modeled as being drawn from a normal distribution with variant-specific mean and variance:

$$y_k \sim \mathcal{N}(\mu_k, \sigma_k^2). \tag{S56}$$

As a result, only two parameters per variant need to be estimated, which amounts to $2K$ parameters in total.

The resulting probabilistic model is depicted graphically in the left half of Fig S1. For completeness, we mention that the same model can also be expressed in terms of the original variables of the multinomial distribution (i.e., the variant fractions $\mathbf{f}$) instead of the transformed variables (i.e., log abundances $\mathbf{y}$) as depicted in the right half of Fig S1. The assumed Gaussian distributions of the components $y_k$ imply a logistic-normal distribution[7] of the fractions $\mathbf{f}$, but with a purely diagonal covariance matrix. The logistic-normal distribution with a general covariance matrix has been promoted by Aitchison for the analysis of compositional data.[8]

##### S3.2.2 Reparametrization trick

The above choices of the functional form for the posterior imply that, when calculating the expectations in (S51), each latent variable $y_k$ should be sampled from a normal distributions with mean $\mu_k$ and variance $\sigma_k^2$. In practice, this can be achieved by generating samples from a standard normal distribution,

$$\epsilon_k \sim \mathcal{N}(0, 1), \tag{S57}$$

and then linearly transforming those as

$$y_k = \mu_k + \sigma_k \epsilon_k. \tag{S58}$$

Rewriting the expectations over the log abundances in (S51) in terms of expectations over the standard normal variables $\boldsymbol{\epsilon}$, we express the ELBO as

$$\mathcal{F}(\theta|\mathbf{n}) = \mathbb{E}_\epsilon\left[\log p(\mathbf{n}|\mathbf{y}(\epsilon))\right] - \mathbb{E}_\epsilon[\log q_\theta(\mathbf{y}(\epsilon))] + \mathbb{E}_\epsilon[\log p(\mathbf{y}(\epsilon))]. \tag{S59}$$

Note that the last two summands are independent of the specific generative model to which the formalism is applied and thus can be calculated before introducing the model.

Starting with the second term, and substituting the transformation (S58) into (S55), we have

$$\log q_\theta(\mathbf{y}(\epsilon)) = -\frac{1}{2}\sum_{k=1}^{K} \epsilon_k^2 - \sum_{k=1}^{K} \log \sigma_k. \tag{S60}$$

Since $\mathbb{E}_\epsilon[\epsilon_k^2] = 1$, we find

$$\mathbb{E}_\epsilon[\log q_\theta(\mathbf{y}(\epsilon))] = -\frac{K}{2} - \sum_{k=1}^{K} \log \sigma_k. \tag{S61}$$

When optimizing the ELBO with respect to $\sigma_k$ the constant term can be dropped.

Moving on to the last term in (S59), we take the prior to be a product of Gaussians with means $\tilde{\mu}_k$ and variances $\tilde{\sigma}_k^2$. Then,

$$\log p(\mathbf{y}(\epsilon)) = -\frac{1}{2}\sum_{k=1}^{K} \frac{1}{\tilde{\sigma}_k^2}(\mu_k + \sigma_k\epsilon_k - \tilde{\mu}_k)^2 - \sum_{k=1}^{K} \log \tilde{\sigma}_k. \tag{S62}$$

Now using $\mathbb{E}_\epsilon[\epsilon_k] = 0$ and $\mathbb{E}_\epsilon[\epsilon_k^2] = 1$ to evaluate the expectation over $\epsilon$, we obtain

$$\mathbb{E}_\epsilon[\log p(\mathbf{y}(\epsilon))] = -\frac{1}{2}\sum_{k=1}^{K} \frac{1}{\tilde{\sigma}_k^2}[(\mu_k - \tilde{\mu}_k)^2 + \sigma_k^2] - \sum_{k=1}^{K} \log \tilde{\sigma}_k. \tag{S63}$$

Here the last term can be dropped when optimizing with respect to the parameters $\mu_k$ and $\sigma_k$ of the posterior.

In summary, retaining only the terms that contain the posterior parameters, we have the ELBO

$$\mathcal{F}(\boldsymbol{\mu}, \boldsymbol{\sigma}|\mathbf{n}) = \mathbb{E}_\epsilon\left[\log p(\mathbf{n}|\mathbf{y}(\epsilon))\right] + \sum_{k=1}^{K} \log \sigma_k - \frac{1}{2}\sum_{k=1}^{K} \frac{1}{\tilde{\sigma}_k^2}[(\mu_k - \tilde{\mu}_k)^2 + \sigma_k^2]. \tag{S64}$$

###### S3.2.3 Uniform prior

In our work, we take uniform priors, which are obtained in the limit $\tilde{\sigma}_k \to \infty$. In this limit, the contribution of the prior to the ELBO in (S64) drops out. The ELBO then simplifies to

$$\mathcal{F}(\boldsymbol{\mu}, \boldsymbol{\sigma}|\mathbf{n}) = \mathbb{E}_\epsilon\left[\mathcal{L}(\mathbf{y}(\boldsymbol{\mu}, \boldsymbol{\sigma}; \epsilon)|\mathbf{n})\right] + \sum_{k=1}^{K} \log \sigma_k \qquad \text{(uniform prior)} \tag{S65}$$

where we also switched to the log likelihood of the generative model, $\mathcal{L}(\mathbf{y}|\mathbf{n})$. This result appears as Eq (20) in the main text.

For the multinomial distribution with *softmax* transformation, which we analyze in the present work, this ELBO is

$$\mathcal{F}(\boldsymbol{\mu}, \boldsymbol{\sigma}|\mathbf{n}) = \mathbb{E}_\epsilon\left[\sum_{k=1}^{K} n_k y_k(\mu_k, \sigma_k; \epsilon_k) - n \log\left(\sum_{k=1}^{K} \mathrm{e}^{y_k(\mu_k, \sigma_k; \epsilon_k)}\right)\right] + \sum_{k=1}^{K} \log \sigma_k. \tag{S66}$$

Substituting (S58) into (S66), and using $\mathbb{E}_\epsilon[\epsilon_k] = 0$ to simplify the first summand, this becomes

$$\mathcal{F}(\boldsymbol{\mu}, \boldsymbol{\sigma}|\mathbf{n}) = \sum_{k=1}^{K} n_k \mu_k - n\mathbb{E}_\epsilon \left[\log\left(\sum_{k=1}^{K} \mathrm{e}^{\mu_k + \sigma_k \epsilon_k}\right)\right] + \sum_{k=1}^{K} \log \sigma_k, \tag{S67}$$

which is Eq (22) of the main text.

#### S3.3 Approximating the expectations and comparison with Dirichlet

The expectation over the standard normal variables $\boldsymbol{\epsilon}$, which appears in the second term of the ELBO (S67), cannot be evaluated analytically and must be approximated. In this subsection, we consider two practical approaches for doing that: one numerical and one analytical. In the numerical strategy, stochastic samples of the random variables are generated, and the expectation of interest is approximated by calculating the mean over the samples (Sec. S3.3.2). This stochastic approximation can be made increasingly more accurate by increasing the number of stochastic samples, which however increases the computational time. In the analytical alternative, instead of maximizing the analytically-intractable ELBO, one maximizes an analytically-tractable lower bound to it (Sec. S3.3.1). The bound that we analyze here is based on the Jensen inequality. By carrying out the maximization analytically, we can show how the widths of the Gaussian distributions relate to the count numbers. However, differently from the stochastic approximation, we have no control over the accuracy of this analytical approximation. As a result, it is not possible to increase the accuracy by increasing the computation time.

##### S3.3.1 Analytical lower bound to the ELBO

For the logarithm function inside the expectation operator in (S67), the Jensen inequality implies that

$$\mathbb{E}_\epsilon \left[\log\left(\sum_{k=1}^{K} \mathrm{e}^{\mu_k + \sigma_k \epsilon_k}\right)\right] \leq \log\left(\sum_{k=1}^{K} \mathbb{E}_\epsilon \left[\mathrm{e}^{\mu_k + \sigma_k \epsilon_k}\right]\right). \tag{S68}$$

By bringing the expectation operator inside the logarithm, it becomes possible to compute the individual expectations analytically. For a standard normal variable $\epsilon_k$, the first moment of the log-normal distribution is

$$\mathbb{E}_\epsilon \left[\mathrm{e}^{\mu_k + \sigma_k \epsilon_k}\right] = \mathrm{e}^{\mu_k + \frac{1}{2}\sigma_k^2}. \tag{S69}$$

Substituting this result back into the ELBO (S67), we find

$$\mathcal{F}(\boldsymbol{\mu}, \boldsymbol{\sigma}|\mathbf{n}) \geq \sum_{k=1}^{K} n_k \mu_k - n \log\left(\sum_{k=1}^{K} \mathrm{e}^{\mu_k + \frac{1}{2}\sigma_k^2}\right) + \sum_{k=1}^{K} \log \sigma_k. \tag{S70}$$

Importantly, for given means, $\boldsymbol{\mu}$, and standard deviations, $\boldsymbol{\sigma}$, the right-hand side of (S70) can be calculated exactly. Since this analytical expression never exceeds the ELBO, it constitutes a lower bound to it. Consequently, instead of maximizing the ELBO, one can infer the variational parameters by maximizing this analytical lower bound.

The gradients of the lower bound with respect to the parameters can be calculated analytically. Equating the derivative with respect to $\mu_k$ to zero leads to

$$\phi_k(\boldsymbol{\mu}^* + \frac{1}{2}\boldsymbol{\sigma}^{*2}) = \frac{n_k}{n}, \tag{S71}$$

where the values of the parameters at the optimum are denoted with the superscript $*$. Similarly, setting the derivative with respect to $\sigma_k$ to zero yields

$$\sigma_k^{*2} = \frac{1}{n_k}. \tag{S72}$$

Thus we managed to obtain an analytical solution for $\sigma_k^*$.

Having determined the standard deviations, we can solve (S71) for the posterior means. Writing the *softmax* function explicitly, and taking the log of both sides, we get

$$\mu_k^* + \frac{1}{2}\sigma_k^{*2} - \log\left(\sum_{i=1}^{K} \mathrm{e}^{\mu_i^* + \frac{1}{2}\sigma_i^{*2}}\right) = \log n_k - \log n. \tag{S73}$$

Since the argument of the *softmax* function is identifiable only up to a global shift, we now discard all variant-independent additive terms and obtain

$$\mu_k^* = \log n_k - \frac{1}{2}(\sigma_k^*)^2 = \log n_k - \frac{1}{2n_k}. \tag{S74}$$

This result constitutes the second half of Eq (28) in the main text.

Thus, for the expected difference of two log abundances we have

$$\mathbb{E}_{\mathrm{LB}}[y_k - y_i] = \mu_k^* - \mu_i^* = \left(\log n_k - \frac{1}{2n_k}\right) - \left(\log n_i - \frac{1}{2n_i}\right). \tag{S75}$$

Similarly, the variance is

$$\mathrm{Var}_{\mathrm{LB}}[y_k - y_i] = \sigma_k^{*2} + \sigma_i^{*2} = \frac{1}{n_k} + \frac{1}{n_i} \qquad (k \neq j), \tag{S76}$$

where we sum the individual variances because the posterior treats the log abundances as independent.

We now use the identity

$$y_i - y_j = \log f_i - \log f_j \tag{S77}$$

to compare the right-hand sides of (S75) and (S76) with the means and variances in (S41), and with their approximations in (S43) and (S44).

Starting with the variance, we see that the right-hand side of (S44) becomes identical to the right-hand side of (S76) if we select $\tilde{\alpha} = 0$ (zero pseudo-counts). Similarly for the mean, the right-hand side of (S43) is identical to the right-hand side of (S75) if $\tilde{\alpha} = 0$. Thus, maximizing the analytical lower-bound to the ELBO essentially recapitulates the Dirichlet analysis with zero pseudo-counts.

**Numerical example** To substantiate the above analytical conclusion, we now analyze the initial counts at $t_0 = 0$ for the illustrative example with four variants considered in the main text. The complete set of counts for exponential growth are

$$\mathbf{N} = \begin{bmatrix} 184 & 245 & 331 & 381 & 416 \\ 97 & 77 & 51 & 16 & 5 \\ 49 & 36 & 13 & 2 & 0 \\ 82 & 17 & 3 & 0 & 0 \end{bmatrix}. \tag{S78}$$

Here we focus on the first column.

With the differences between variants $i$ and $j$ ordered in a matrix, the means and variances calculated acording to (S75) and (S76) are, respectively,

$$\left[\mathbb{E}_{\mathrm{LB}}[y_{i0} - y_{j0}]\right] = \left[\mu_i^* - \mu_j^*\right] = \begin{bmatrix} 0 & 0.643 & 1.331 & 0.812 \\ -0.643 & 0 & 0.688 & 0.169 \\ -1.331 & -0.688 & 0 & -0.519 \\ -0.812 & -0.169 & 0.519 & 0 \end{bmatrix} \tag{S79}$$

and

$$\left[\mathrm{Var}_{\mathrm{LB}}[y_{i0} - y_{j0}]\right] = \left[\sigma_i^{*2} + \sigma_j^{*2}\right] = \begin{bmatrix} & 0.016 & 0.026 & 0.018 \\ 0.016 & & 0.031 & 0.023 \\ 0.026 & 0.031 & & 0.033 \\ 0.018 & 0.023 & 0.033 & \end{bmatrix}. \tag{S80}$$

(The index $i$ runs along the rows and $j$ along the columns.) Note that the means in (S79) change sign upon flipping the order of $i$ and $j$, while the variances in (S80) remain unchanged. The corresponding matrices are, therefore, antisymmetric and symmetric, respectively.

With $\tilde{\alpha} = 0$, according to (S37) the posterior Dirichlet parameters are directly equal to the variant counts. The means and variances calculated from (S41), i.e., using the exact digamma and trigamma functions, are

$$\big[\mathbb{E}_{\mathrm{Dir}}[\log f_{i0} - \log f_{j0}]\big] = \begin{bmatrix} 0 & 0.643 & 1.331 & 0.812 \\ -0.643 & 0 & 0.688 & 0.169 \\ -1.331 & -0.688 & 0 & -0.519 \\ -0.812 & -0.169 & 0.519 & 0 \end{bmatrix} \qquad (\tilde{\alpha} = 0) \tag{S81}$$

$$\big[\mathrm{Var}_{\mathrm{Dir}}[\log f_{i0} - \log f_{j0}]\big] = \begin{bmatrix} & 0.016 & 0.026 & 0.018 \\ 0.016 & & 0.031 & 0.023 \\ 0.026 & 0.031 & & 0.033 \\ 0.018 & 0.023 & 0.033 & \end{bmatrix} \qquad (\tilde{\alpha} = 0). \tag{S82}$$

As expected from the analytical derivation, these values are identical to the ones obtained by minimizing the lower bound to the ELBO because the initial counts in (S78) are relatively large and so the functions $\psi_0(\cdot)$ and $\psi_1(\cdot)$ are well approximated by the first terms in their asymptotic expansions.

To see the quantitative effect of the prior, we repeated the Dirichlet calculation using a uniform prior ($\tilde{\alpha} = 1$). The resulting means and variances in this case are

$$\big[\mathbb{E}_{\mathrm{Dir}}[\log f_{i0} - \log f_{j0}]\big] = \begin{bmatrix} 0 & 0.638 & 1.316 & 0.805 \\ -0.638 & 0 & 0.678 & 0.167 \\ -1.316 & -0.678 & 0 & -0.511 \\ -0.805 & -0.167 & 0.511 & 0 \end{bmatrix} \qquad (\tilde{\alpha} = 1) \tag{S83}$$

$$\big[\mathrm{Var}_{\mathrm{Dir}}[\log f_{i0} - \log f_{j0}]\big] = \begin{bmatrix} & 0.016 & 0.026 & 0.018 \\ 0.016 & & 0.030 & 0.022 \\ 0.026 & 0.030 & & 0.032 \\ 0.018 & 0.022 & 0.032 & \end{bmatrix} \qquad (\tilde{\alpha} = 1). \tag{S84}$$

Not surprisingly, the prior has a quantitative effect on the inferred values, although this effect is negligibly small for the calculated variances. It is also small for the means, since in the present case of about fifty and more counts per variant adding $\tilde{\alpha} = 0$ or $\tilde{\alpha} = 1$ pseudo-counts does not make much of a difference. The effect is expected to be more pronounced in the case of smaller count numbers.

##### S3.3.2 Monte Carlo approximation

If maximizing the lower bound to the ELBO recapitulates the Dirichlet inference, what does maximizing the actual ELBO achieve? To address this question, we turn to the stochastic approximation of the expectation term in (S67).

From the perspective of sampling, where one actually generates samples of the random variables, this expectation corresponds to the average over infinitely many independent random realizations. Specifically, if we denote one random realization by $\boldsymbol{\epsilon}^{[m]}$, then

$$\mathbb{E}_{\epsilon}\left[\log\left(\sum_{k=1}^{K} \mathrm{e}^{\mu_k + \sigma_k \epsilon_k}\right)\right] = \lim_{M\to\infty} \frac{1}{M} \sum_{m=1}^{M} \log\left(\sum_{k=1}^{K} \mathrm{e}^{\mu_k + \sigma_k \epsilon_k^{[m]}}\right). \tag{S85}$$

By calculating the sample average over a finite number of realizations, $M$, we obtain an approximation to the exact expectation. Because the sample mean over another batch of $M$ random realizations will be different, in general, the obtained approximate value will change stochastically every time it is calculated. As a result, during the maximization of the ELBO, we actually work with its stochastic approximation

$$\mathcal{F}_M(\boldsymbol{\mu}, \boldsymbol{\sigma}|\mathbf{n}) = \sum_{k=1}^{K} n_k \mu_k - n \frac{1}{M} \sum_{m=1}^{M} \log\left(\sum_{k=1}^{K} \mathrm{e}^{\mu_k + \sigma_k \epsilon_k^{[m]}}\right) + \sum_{k=1}^{K} \log \sigma_k. \tag{S86}$$

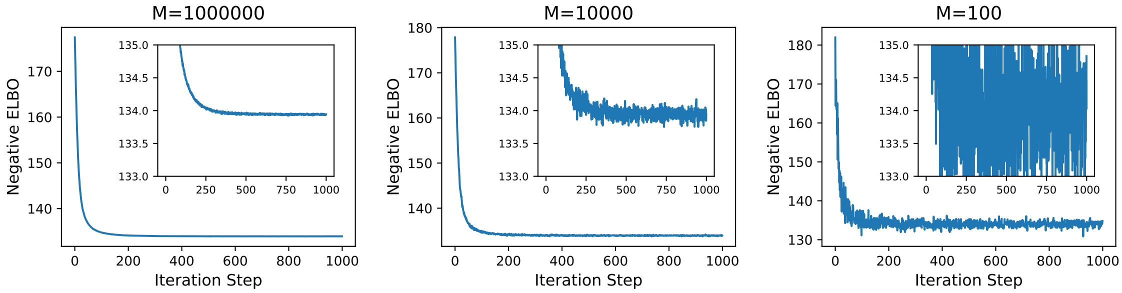


Figure S2: Stochastic fluctuations of the ELBO increase for decreasing sample size $M$. The inset shows the same values as the main plot but restricted to a fixed interval along the vertical axis. (The gradient-descent optimization was initialized with means equal to zero and standard deviations equal to one.)

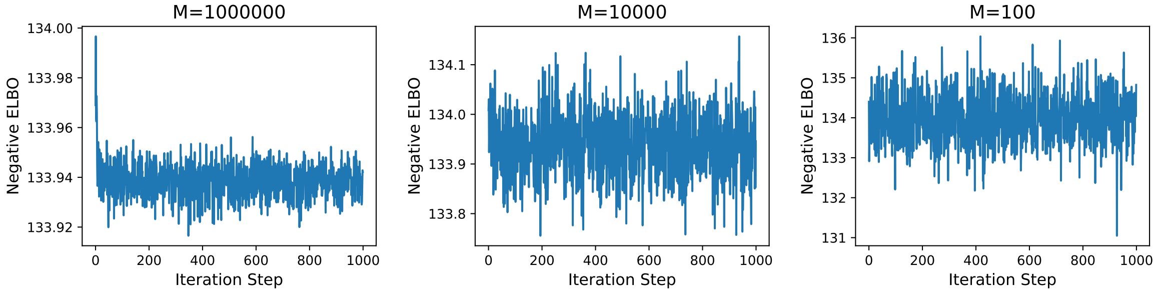


Figure S3: Same as Fig S2 but with variational parameters initialized as $\mu_k = \log n_k - 1/2n_k$ and $\log \sigma_k = -(\log n_k)/2$, which maximize the Jensen lower bound to the ELBO.

Naturally, the gradients of this stochastic $\mathcal{F}_M$ with respect to the parameters $\boldsymbol{\mu}$ and $\boldsymbol{\sigma}$, which are accumulated automatically during the numerical integration of the dynamics, will also be stochastic approximations to the true gradients, and so will continue to fluctuate even at the optimum.

**Numerical example** To illustrate the effect of the sample size $M$, we again analyze the initial counts in the first column of (S78) for different choices of $M$. In Fig S2 we show the negative ELBO as a function of the gradient-descent iteration steps for $M = 10^6, 10^4$ and $10^2$. In each case, the minimization is carried out for 1000 iteration steps. The insets zoom into a fixed range of the vertical axis.

When the sample size is relatively small ($M = 100$), after the initial steep decrease, the stochastic approximation to the ELBO fluctuates wildly from one iteration step to the next. The fluctuations decrease when $M$ is increased to $10^4$ and $10^6$, since the Monte Carlo approximations to the expectation become increasingly more accurate.

The initial rapid decay of the negative ELBO observed in Fig S2 indicates that the gradient-descent optimization is initialized with values that are relatively far from the optimum values. (All means were set to zero and all standard deviations to one.) Starting with values that are closer to the optimal ones would reduce the number of steps needed to reach the optimum.

As better initial values, we use the means and standard deviations in (S74) and (S72), obtained by maximizing the Jensen lower bound to the ELBO. The resulting optimization runs, with other settings left unchanged compared to Fig S2, are shown in Fig S3. In this case, the initial decay of the negative ELBO is visible only for $M = 10^6$ samples; for $M = 10^4$ and $M = 10^2$, the difference between the initial and optimal ELBO is smaller than the amplitude of the stochastic fluctuations.

Now that the initial values are close to the optimum, we reduce the fixed number of optimization steps from 1000 to 500.

Because of the persistent stochastic fluctuations, we assess convergence by visually inspecting the negative ELBO. Since the variational parameters continue to fluctuate throughout the optimization, we average over the last 100 steps when reporting final estimates. For the standard deviations, whose logarithms are actually used as optimization parameters, we calculate both the mean and variance of $\log \sigma_k$ over the last 100 iteration steps, and use these to estimate the mean of $\sigma_k$, analogously to (S69).

When $M = 10^6$, the parameter values at the last optimization step and the averages over the last 100 steps were identical to within at least three digits after the decimal point. The means and variances we obtained in this high-precision case were

$$\left[\mathbb{E}_{\mathrm{VI}}[y_{i0} - y_{j0}]\right] = \left[\mu_i^{\mathrm{VI}} - \mu_j^{\mathrm{VI}}\right] = \begin{bmatrix} 0 & 0.643 & 1.331 & 0.812 \\ -0.643 & 0 & 0.688 & 0.169 \\ -1.331 & -0.688 & 0 & -0.519 \\ -0.812 & -0.169 & 0.519 & 0 \end{bmatrix} \qquad (M = 10^6) \tag{S87}$$

and

$$\left[\mathrm{Var}_{\mathrm{VI}}[y_{i0} - y_{j0}]\right] = \left[(\sigma_i^{\mathrm{VI}})^2 + (\sigma_j^{\mathrm{VI}})^2\right] = \begin{bmatrix} & 0.026 & 0.037 & 0.028 \\ 0.026 & & 0.040 & 0.032 \\ 0.037 & 0.040 & & 0.042 \\ 0.028 & 0.032 & 0.042 & \end{bmatrix} \qquad (M = 10^6). \tag{S88}$$

(The superscript VI denotes *variational-inference* values obtained by maximizing the actual ELBO.) The means are identical to the ones we obtained before by maximizing the lower bound to the ELBO and from the Dirichlet analysis without pseudo-counts (i.e., $\tilde{\alpha} = 0$). In contrast, the current variances are larger, hence the uncertainties assigned to the inferred parameters by maximizing the actual ELBO—and thus minimizing the KL divergence—are larger than those of the Dirichlet analysis.

To assess the effect of reducing the sample size, we repeated the same analysis with $M = 100$. The resulting means and variances (averaged over the last 100 steps) are

$$\left[\mathbb{E}_{\mathrm{VI}}[y_{i0} - y_{j0}]\right] = \left[\mu_i^{\mathrm{VI}} - \mu_j^{\mathrm{VI}}\right] = \begin{bmatrix} 0 & 0.642 & 1.331 & 0.810 \\ -0.642 & 0 & 0.690 & 0.169 \\ -1.331 & -0.690 & 0 & -0.521 \\ -0.810 & -0.169 & 0.521 & 0 \end{bmatrix} \qquad (M = 100) \tag{S89}$$

and

$$\left[\mathrm{Var}_{\mathrm{VI}}[y_{i0} - y_{j0}]\right] = \left[(\sigma_i^{\mathrm{VI}})^2 + (\sigma_j^{\mathrm{VI}})^2\right] = \begin{bmatrix} & 0.026 & 0.036 & 0.027 \\ 0.026 & & 0.040 & 0.032 \\ 0.036 & 0.040 & & 0.042 \\ 0.027 & 0.032 & 0.042 & \end{bmatrix} \qquad (M = 100). \tag{S90}$$

Although these numerical values are different than the high-precision values above, it is clear that samples that are much smaller than $M = 10^6$ can be used, depending on the desired precision.

So far we focused on the difference between two variants, as given in (S77). Of ultimate interest, however, are the absolute values of the individual means and variances, $\mu_k$ and $\sigma_k^2$, and not only their differences or sums. Next, we check how the inferred values of $\boldsymbol{\mu}^{\mathrm{VI}}$ and $\boldsymbol{\sigma}^{\mathrm{VI}}$ depend on the Monte Carlo sample size.

For $M = 10^3$, we now repeat the described optimization $R = 100$ times, and report the mean values and the standard deviations over these repetitions:

$$\begin{aligned} \boldsymbol{\mu}_{1000}^{\mathrm{VI}} &= \left[-0.8036 \pm 0.0002 \quad -1.4468 \pm 0.0003 \quad -2.1350 \pm 0.0003 \quad -1.6159 \pm 0.0003\right] \\ \boldsymbol{\sigma}_{1000}^{\mathrm{VI}} &= \left[0.1001 \pm 0.0027 \quad 0.1169 \pm 0.0019 \quad 0.1525 \pm 0.0020 \quad 0.1234 \pm 0.0020\right]. \end{aligned} \tag{S91}$$

The small variation in the means over the $R = 100$ repetitions, indicates that these can be calculated rather accurately even with a smaller $M$.

We further reduced the number of random samples per variant by one order of magnitude to $M = 100$, and increased the number of repeated minimizations to $R = 1000$ for better statistics. The result was

$$\begin{aligned}\boldsymbol{\mu}_{100}^{\mathrm{VI}} &= \begin{bmatrix}-0.8036 \pm 0.0006 & -1.4468 \pm 0.0009 & -2.1350 \pm 0.0011 & -1.6159 \pm 0.0009\end{bmatrix} \\ \boldsymbol{\sigma}_{100}^{\mathrm{VI}} &= \begin{bmatrix}0.1007 \pm 0.0080 & 0.1176 \pm 0.0063 & 0.1530 \pm 0.0055 & 0.1243 \pm 0.0060\end{bmatrix}.\end{aligned} \tag{S92}$$

The uncertainties of the inferred values increased roughly by a factor of three, which is the square root of 10. Going a step further and using $M = 10$, we get

$$\begin{aligned}\boldsymbol{\mu}_{10}^{\mathrm{VI}} &= \begin{bmatrix}-0.8034 \pm 0.0022 & -1.4471 \pm 0.0033 & -2.1351 \pm 0.0040 & -1.6158 \pm 0.0034\end{bmatrix} \\ \boldsymbol{\sigma}_{10}^{\mathrm{VI}} &= \begin{bmatrix}0.1046 \pm 0.0193 & 0.1198 \pm 0.0158 & 0.1547 \pm 0.0155 & 0.1276 \pm 0.0151\end{bmatrix}\end{aligned} \tag{S93}$$

and the uncertainties increased by another factor of three.

This numerical exploration shows that, depending on the desired precision, the expectations over the standard normal variables can be approximated by using as little as ten random samples per variant.

##### S3.3.3 Further comparison with the Dirichlet analysis

The described variational inference yields the means and variances of the log abundances $y_k$:

$$\mathbb{E}[y_k] = \mu_k, \qquad \mathrm{Var}[y_k] = \sigma_k^2. \tag{S94}$$

Given this information, we now want to determine the means and variances of the fractions $f_k$. Recall that the relationship between these two sets of random variables is given by the *softmax* transformation (S13), which we write here again but without showing the time index:

$$f_k = \phi_k(\mathbf{y}) = \frac{\mathrm{e}^{y_k}}{\sum_{i=1}^{K} \mathrm{e}^{y_i}}. \tag{S95}$$

Because the exact means and variances of $f_k$ cannot be obtained analytically from those of $y_k$, here we derive approximate expressions using the so-called *delta method*, which is based on the Taylor expansion of the *softmax* transformation.

Expanding the *softmax* function (S95) around the mean of its argument, and truncating the expansion at the first-order term, we get

$$f_k \approx \phi_k(\boldsymbol{\mu}) + \sum_{i=1}^{K} \left.\frac{\partial \phi_k}{\partial y_i}\right|_{\mathbf{y}=\boldsymbol{\mu}} (y_i - \mu_i). \tag{S96}$$

Taking the expectation of both sides, we find that, to this order, the means of the fractions are obtained by *softmax* transforming the means of the original variables:

$$\mathbb{E}[f_k] \approx \phi_k(\boldsymbol{\mu}). \tag{S97}$$

Moving on to the variances, we start with the definition of variance and then substitute the approximate mean from (S97) to arrive at

$$\mathrm{Var}[f_k] \approx \mathbb{E}[(f_k - \phi_k(\boldsymbol{\mu}))^2] \approx \mathbb{E}\left[\left(\sum_{i=1}^{K} \left.\frac{\partial \phi_k}{\partial y_i}\right|_{\mathbf{y}=\boldsymbol{\mu}} (y_i - \mu_i)\right)^2\right]. \tag{S98}$$

where the truncated Taylor expansion in (S96) was used in the second approximation. Because the log abundances are treated as independent in the mean field approximation, only the "diagonal" terms of the square on the right-hand side of (S98) will survive the expectation operation. Thus,

$$\mathrm{Var}[f_k] \approx \sum_{i=1}^{K} \left(\left.\frac{\partial \phi_k}{\partial y_i}\right|_{\mathbf{y}=\boldsymbol{\mu}}\right)^2 \mathbb{E}[(y_i - \mu_i)^2] = \sum_{i=1}^{K} \left(\left.\frac{\partial \phi_k}{\partial y_i}\right|_{\mathbf{y}=\boldsymbol{\mu}}\right)^2 \sigma_i^2, \tag{S99}$$

Table S1: Posterior means and standard deviations of the variant fractions from the exact (Dirichlet) and variational (ELBO) Bayesian analysis of the counts at the initial time $t_0$.

| | $\mathbb{E}[\mathbf{f}_0]$ | $\text{Std}[\mathbf{f}_0]$ |
|---|---|---|
| Dirichlet ($\tilde{\alpha} = 1$) | $[0.445\ \ 0.236\ \ 0.120\ \ 0.200]$ | $[0.024\ \ 0.021\ \ 0.016\ \ 0.020]$ |
| ELBO (lower bound) | $[0.448\ \ 0.235\ \ 0.118\ \ 0.199]$ | $[0.025\ \ 0.021\ \ 0.016\ \ 0.020]$ |
| ELBO (stochastic $M = 10^6$) | $[0.448\ \ 0.235\ \ 0.118\ \ 0.199]$ | $[0.031\ \ 0.024\ \ 0.017\ \ 0.023]$ |
| True $\mathbf{f}_0$ | $[0.385\ \ 0.256\ \ 0.132\ \ 0.227]$ | - |

where in the last equality we recognized the variance of $y_i$ to be $\sigma_i^2$. Thus, to this order, the variance of $f_k$ is obtained as the weighted sum of the variances of the original variables, with the weights given by the squares of the components of the gradient of the transformation at $\boldsymbol{\mu}$.

From (S95), we find the gradient to be

$$\frac{\partial \phi_k}{\partial y_i} = \begin{cases} \phi_k(\mathbf{y})[1 - \phi_k(\mathbf{y})], & i = k \\ -\phi_k(\mathbf{y})\phi_i(\mathbf{y}), & i \neq k. \end{cases} \tag{S100}$$

Evaluating at $\boldsymbol{\mu}$, and substituting on the right-hand side of (S99), we get

$$\text{Var}[f_k] \approx [\phi_k(\boldsymbol{\mu})]^2 \left\{ [1 - 2\phi_k(\boldsymbol{\mu})]\sigma_k^2 + \sum_{i=1}^{K} [\phi_i(\boldsymbol{\mu})]^2 \sigma_i^2 \right\}, \tag{S101}$$

and taking the square root of both sides,

$$\text{Std}[f_k] \approx \phi_k(\boldsymbol{\mu})\sigma_k \left\{ 1 - 2\phi_k(\boldsymbol{\mu}) + \frac{1}{\sigma_k^2} \sum_{i=1}^{K} \sigma_i^2 [\phi_i(\boldsymbol{\mu})]^2 \right\}^{\frac{1}{2}}. \tag{S102}$$

The means and standard deviations of the fractions $f_k$ calculated from these approximations are given in the second and third rows of Table S1. The values in the second row are calculated from $\boldsymbol{\mu}^*$ and $\boldsymbol{\sigma}^*$, obtained by maximizing the lower bound to the ELBO, and those in the third row from $\boldsymbol{\mu}^{\text{VI}}$ and $\boldsymbol{\sigma}^{\text{VI}}$, obtained by maximizing the stochastic approximation to the ELBO with $M = 10^6$ random samples. These statistics can now be compared directly with those of the Dirichlet analysis in the first row of the table, obtained for a uniform prior ($\tilde{\alpha} = 1$). Despite the approximate nature of (S97), the means of the fractions are in excellent agreement. For the standard deviations, maximizing the lower bound to the ELBO recapitulates the Dirichlet analysis, while maximizing the ELBO yields larger uncertainties.

For comparison, the true fractions used to generate the counts data, are given in the last row of the table. Since our inference is based on a single realization of the multinomial random variable, we do not expect to recover the true values of the fractions. Nonetheless, these fall within three posterior standard deviations of the posterior means.

### S4 Simultaneous treatment of the counting noise at all time points

In Secs. S2 and S3, each sequencing time point was analyzed in isolation from the others. In this section, we give the log likelihood and ELBO that combine the counts across all time points.

#### S4.1 Maximum likelihood

If the counting noise at each sequencing time point is independent from the noise at the other time points, the single-time log likelihoods in (S24) can be added together to produce the combined log likelihood

$$\mathcal{L}(\mathbf{Y}|\mathbf{N}) = \sum_{\tau=0}^{T-1} \left[ \sum_{k=1}^{K} n_{k\tau} y_{k\tau} - n_\tau \log \left( \sum_{k=1}^{K} \mathrm{e}^{y_{k\tau}} \right) \right], \tag{S103}$$

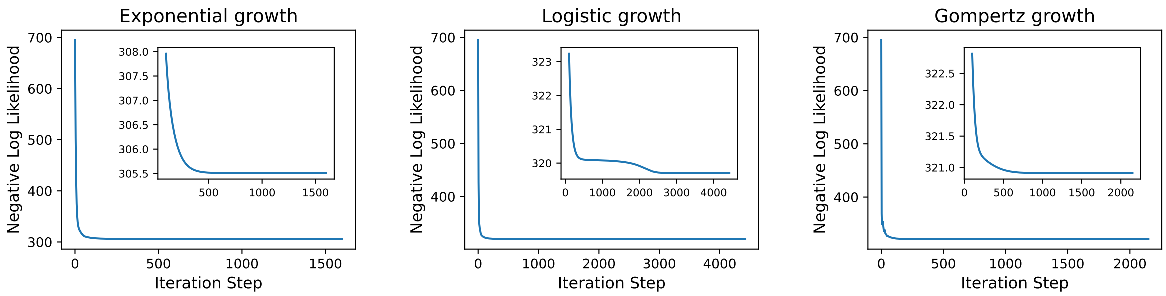


Figure S4: Negative of the log likelihood as a function of the gradient descent iteration steps for the example with $K = 4$ variants. Insets show values starting from iteration step 100. (Learning rate: 0.01.)

which relates the log abundances at all times, $\mathbf{Y} = [y_{k\tau}]$, to the counts at all times, $\mathbf{N} = [n_{k\tau}]$. ($\mathbf{Y}$ and $\mathbf{N}$ are $K \times T$ matrices with rows corresponding to the variants and columns corresponding to the different sequencing time points.)

Note that the sum over the time points in (S103) does not actually achieve the desired collective analysis of all counts. Indeed, maximizing this log likelihood with respect to the log abundances $y_{k\tau}$ would recover the result of Sec. S2.1.2, where each time point was analyzed separately. The desired integration is achieved through the growth model, which expresses the log abundances at different time points in terms of the time-independent growth parameters:

$$y_{k\tau} = y_k(\boldsymbol{\lambda}, \boldsymbol{\delta}; t_\tau). \tag{S104}$$

Substituting into (S103) yields

$$\mathcal{L}(\boldsymbol{\lambda}, \boldsymbol{\delta}|\mathbf{N}) = \sum_{\tau=0}^{T-1} \sum_{k=1}^{K} n_{k\tau} y_k(\boldsymbol{\lambda}, \boldsymbol{\delta}; t_\tau) - \sum_{\tau=0}^{T-1} n_\tau \log \left( \sum_{k=1}^{K} \mathrm{e}^{y_k(\boldsymbol{\lambda}, \boldsymbol{\delta}; t_\tau)} \right), \tag{S105}$$

which can now be maximized directly with respect to the growth parameters.

We minimized the negative of the log likelihood in (S105) using the counts data given in the main text, and reproduced here in (S78) for exponential growth. The gradual decrease of the loss function during the minimization is shown in Fig S4 for the three growth models. The optimization was stopped when the log likelihood changed by less than $10^{-12}$ and the maximum change of the values of the parameters was less than $10^{-10}$ from one iteration to the next. The number of iteration steps in Fig S4 differs across the three growth models since the stopping criterion was reached at different points. The variant fractions calculated from the estimated maximum-likelihood growth parameters are given in Fig 4 of the main text.

#### S4.2 Variational inference

Now we promote the variant-specific parameters of the growth model—growth rates $\boldsymbol{\lambda}$ and initial log abundances $\boldsymbol{\delta}$—to normal random variables. The means and standard deviations of their normal distributions become the parameters that need to be inferred from the data. Analogously to maximizing the likelihood, here we maximize the ELBO

$$\mathcal{F}(\boldsymbol{\mu}^\lambda, \boldsymbol{\sigma}^\lambda, \boldsymbol{\mu}^\delta, \boldsymbol{\sigma}^\delta|\mathbf{N}) = \sum_{\tau=0}^{T-1} \sum_{k=1}^{K} n_{k\tau} \mathbb{E}_\epsilon[y_{k\tau}] - \sum_{\tau=0}^{T-1} n_\tau \mathbb{E}_\epsilon \left[ \log \left( \sum_{k=1}^{K} \mathrm{e}^{y_{k\tau}} \right) \right] + \sum_{k=1}^{K} \log \sigma_k^\lambda + \sum_{k=1}^{K} \log \sigma_k^\delta, \tag{S106}$$

where

$$y_{k\tau} = y_k(\boldsymbol{\lambda}(\boldsymbol{\mu}^\lambda, \boldsymbol{\sigma}^\lambda; \boldsymbol{\epsilon}^\lambda), \boldsymbol{\delta}(\boldsymbol{\mu}^\delta, \boldsymbol{\sigma}^\delta; \boldsymbol{\epsilon}^\delta); t_\tau) \tag{S107}$$

and

$$\boldsymbol{\lambda}(\boldsymbol{\mu}^\lambda, \boldsymbol{\sigma}^\lambda; \boldsymbol{\epsilon}^\lambda) = \boldsymbol{\mu}^\lambda + \boldsymbol{\sigma}^\lambda \odot \boldsymbol{\epsilon}^\lambda, \qquad \boldsymbol{\delta}(\boldsymbol{\mu}^\delta, \boldsymbol{\sigma}^\delta; \boldsymbol{\epsilon}^\delta) = \boldsymbol{\mu}^\delta + \boldsymbol{\sigma}^\delta \odot \boldsymbol{\epsilon}^\delta. \tag{S108}$$

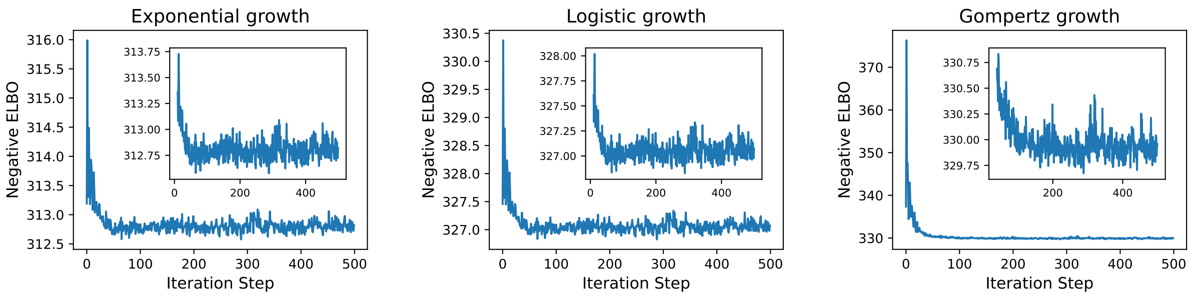


Figure S5: Negative ELBO from Monte Carlo approximation with $M = 100$, for a fixed duration of 500 iteration steps with learning rate of 0.03. Insets start from step 10 for exponential and logistic growth, and from step 40 for Gompertz growth.

The expectation $\mathbb{E}_\epsilon[\cdot]$ in the ELBO is now over the independent standard normal variables $\boldsymbol{\epsilon}^\lambda$ and $\boldsymbol{\epsilon}^\delta$.

We approximate these expectations stochastically, as explained earlier in Sec. S3.3.2, by drawing random samples of the standard normal variables and calculating the sample mean. For each variant, we generate $M$ random pairs $(\epsilon_k^\lambda, \epsilon_k^\delta)$ and use them to obtain $M$ realizations of the growth parameters $(\lambda_k, \delta_k)$ according to (S108). The parameters with the same sample index are combined across the variants to integrate the growth dynamics either analytically (exponential growth) or numerically (logistic and Gompertz growth). The expectations in the ELBO are then approximated by averaging over these $M$ realizations of the growth trajectories of all variants.

##### S4.2.1 Synthetic data with four variants

We carried out the numerical minimization of the negative ELBO for the illustrative counts data in the main text. The means $\boldsymbol{\mu}^\lambda$ and $\boldsymbol{\mu}^\delta$ at the start of the optimization were set equal to the values $\boldsymbol{\lambda}^{\mathrm{ML}}$ and $\boldsymbol{\delta}^{\mathrm{ML}}$ that maximized the log likelihood. The logarithms of the standard deviations were initialized as

$$\log \sigma_k^\lambda = -\frac{1}{2} \log \left( \sum_{\tau=0}^{T-1} t_\tau^2 n_{k\tau} \right), \qquad \log \sigma_k^\delta = -\frac{1}{2} \log \left( \sum_{\tau=0}^{T-1} n_{k\tau} \right). \tag{S109}$$

This choice is motivated by the Jensen lower bound to the ELBO (S106) for exponential growth, presented in Sec 3.3.1 of the main text.

The decrease of the negative ELBO during 500 optimization steps is shown in Fig S5 for the three growth models. The first two models appear to converge already within the first 100 steps, after which the stochastic fluctuations about the mean persist. Convergence takes about 200 steps in the case of Gompertz growth.

The final means and standard deviations were obtained by averaging the over the last 100 iteration steps, as explained earlier. These averaged values were visualized as Gaussians in Fig 5 of the main text. The means and standard deviations of the growth rates were given in Table 4 of the main text. In Table S2, we give the means and standard deviations of the initial log abundances across the three growth models.

##### S4.2.2 Synthetic data with one hundred variants

Here we show supporting figures for the example with $K = 100$ variants under logistic growth, whose posterior distributions are plotted in Fig 8A of the main text. In the left panel of Fig S6, we show a histogram of the variants with non-zero counts across all five time points. Note that at least two non-zero time points are needed to estimate variant growth rate and initial abundance.

The decay of the negative log likelihood during the maximum-likelihood optimization is shown in middle panel of Fig S6. The stopping criterion in this case was that the log likelihood and the individual parameters change by less than $10^{-4}$ and $10^{-4}$, respectively, within a single iteration step with learning rate of 0.01.

Table S2: Posterior means and standard deviations of the initial log abundances obtained by maximizing the all-times ELBO for the example with $K = 4$ variants. The corresponding growth rates are given in Table 4 of the main text.

| | $\boldsymbol{\mu}^\delta$ (in gauge $\sum_k e^{\mu_k} = 1$) | | | | $\boldsymbol{\sigma}^\delta$ (uncertainties) | | | |
|---|---|---|---|---|---|---|---|---|
| Exponential | $-0.843$ | $-1.427$ | $-2.040$ | $-1.612$ | 0.063 | 0.072 | 0.104 | 0.109 |
| Logistic | $-0.832$ | $-1.434$ | $-2.068$ | $-1.610$ | 0.063 | 0.070 | 0.102 | 0.109 |
| Gompertz | $-0.830$ | $-1.433$ | $-2.039$ | $-1.633$ | 0.068 | 0.069 | 0.100 | 0.113 |
| True $\boldsymbol{\delta}$ | $-0.955$ | $-1.363$ | $-2.026$ | $-1.481$ | - | | | |

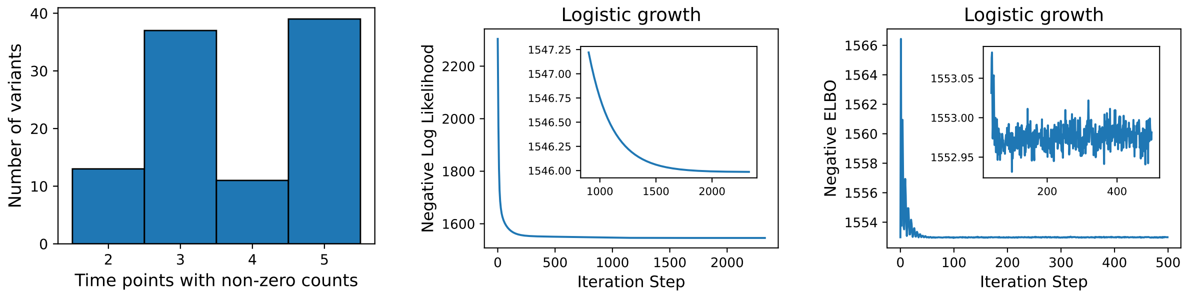


Figure S6: Details about the numerical example with $K = 100$ variants. Left: Histogram of the variants having the specified number of non-zero counts at the five sequencing time points. Middle: Optimization of the log likelihood with learning rate of 0.01. Right: Optimization of the ELBO for 500 steps with $M = 100$ and learning rate 0.03.

The estimates obtained by minimizing the log likelihood were used as starting values for the variational means in the optimization of the ELBO. The values of the negative ELBO during minimization for 500 steps are shown in the right panel of Fig S6. The values of the variational parameters averaged over the last 100 iteration steps were used to generate Fig 8 of the main text.

##### S4.2.3 Synthetic data with one thousand variants

Information about the analysis with $K = 1000$ variants, which is mentioned in the main text, is summarized in Fig S7.

##### S4.2.4 Experimental data with five thousand variants

Although not mentioned in the main text, we also analyzed actual experimental data from Ghosh et al.[4] on the growth of yeast variants, generously made available at `https://github.com/omghosh/limiting-functions`.

From the complete dataset, we arbitrarily selected replicate `R2` of `Batch3`, grown in `Suc` at dilution frequency of `1Day`. The four time points for this condition, together with the starting time point that is common to all conditions, amount to five sequencing time points, separated by 24 hours.

Information about the analysis of these variants is provided in Fig S8. From the start we dropped ten of the variants, since they either had not been observed at any of the five sequencing time points or had been observed only once (orange bars). For the remaining 5017 variants, we first performed maximum-likelihood estimation, followed by variational inference. Since we used a model of exponential growth, both the growth rates and the initial log abundances of the variants can be globally shifted to any desired gauge. We constrained the sum of the growth rates to zero and the exponential sum of the initial log abundances to one.

The correlations between the means and precisions (i.e., inverse variance) of the growth parameters are shown in the right panel of Fig S8. These plots reveal the presence of two groups of variants that have

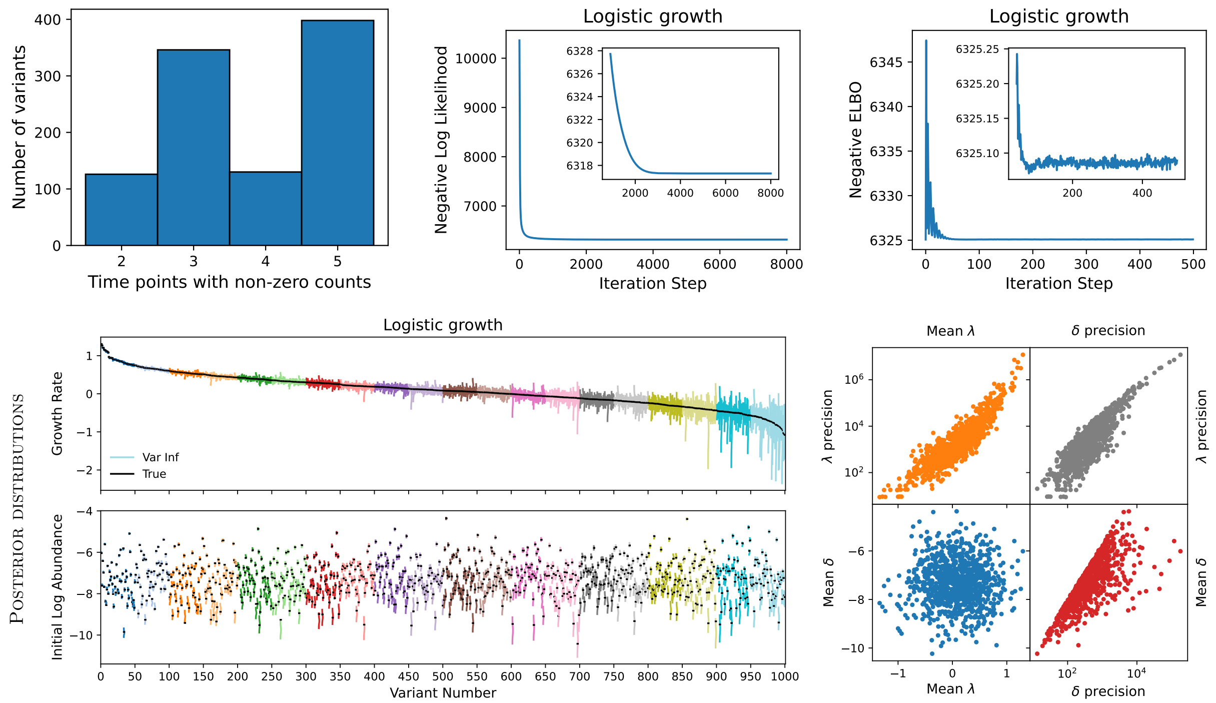


Figure S7: Details about the numerical example with $K = 1000$ variants. Top-row panels are comparable to those in Fig S6 for 100 variants, while bottom-row panels are analogous to those in Fig 7 of the main text.

very high initial abundance (Mean $\delta$), but intermediate and low growth rates (Mean $\lambda$). (These two groups may have been spiked into the pool of all other variants. One of them could presumably be the wild type.) Because of their high initial abundance, the variants in these two groups also have relatively low growth-rate uncertainties (i.e., high $\lambda$ precisions).

Additional figures for these data can be found at `https://github.com/dzsezer/growth_rates_from_counts/`.

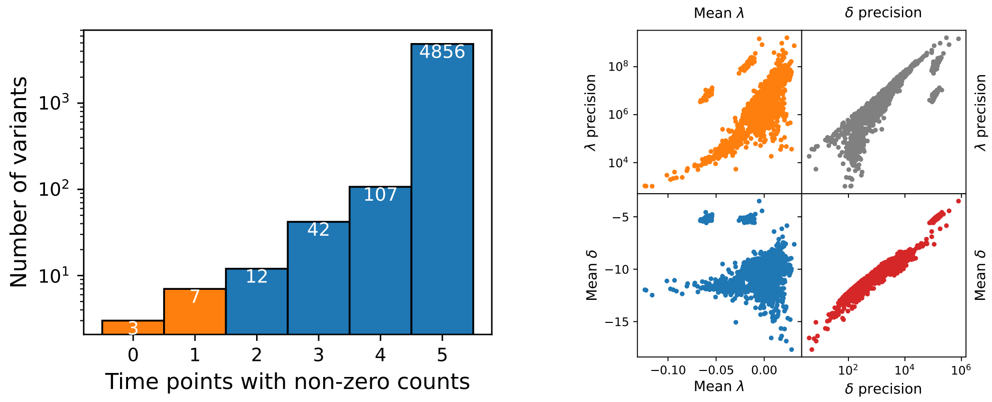


Figure S8: Details about the experimental data with five thousand variants.

[8] John Aitchison. *The Statistical Analysis of Compositional Data*. Monographs on Statistics and Applied Probability. Chapman and Hall, London, 1986. ISBN 0412280604.
